# Rab12 is a regulator of mitophagy and mitochondrial homeostasis

**DOI:** 10.64898/2026.03.29.715103

**Authors:** Tara Richbourg, Alex George, Anas Bitar, Ian T. Ryde, Casey Farrell, Tuyana Malankhanova, Jennifer Liu, Silas A. Buck, Ivana Barraza, So Young Kim, Xiaoyan Nie, Andrew B. West, Joel N. Meyer, Laurie H. Sanders

## Abstract

Rab GTPases orchestrate vesicular trafficking, but their contributions to mitochondrial quality control are not fully defined, despite links to multiple mitochondria-related human diseases. We conducted a family-wide siRNA-based screen using mt-mKeima/YFP–Parkin HeLa cells to identify regulators of depolarization-induced mitophagy. The screen identified several candidate Rabs, and follow-up studies validated Rab12 as a negative regulator of mitophagy. Rab12 knockdown or knockout augments clearance of damaged mitochondria basally and/or after FCCP-induced depolarization, with findings reproduced across distinct cell types. Rab12 depletion increased mitochondrial content, lowered mitochondrial membrane potential, and reduced mitochondrial DNA damage, without detectable changes in overall cellular bioenergetic capacity. Together, these results indicate that Rab12 restrains mitophagic engagement and its loss permits accumulation of lower-functioning mitochondria that are hypersensitive to mitophagy-inducing stress. Rab12 thus emerges as a novel effector linking vesicular trafficking machinery and mitochondrial homeostasis, with potential implications for neurodegenerative disorders and other Rab-associated diseases.

## INTRODUCTION

Ras-associated binding (Rab) proteins comprise a large family of small GTPases that act as master regulators of vesicular trafficking in eukaryotic cells.^1,2^ By cycling between GDP-bound (inactive) and GTP-bound (active) states, Rabs associate with intracellular membranes to regulate vesicle budding, transport, tethering, and fusion.^2,3^ Because these functions underpin intracellular membrane trafficking and cellular homeostasis, dysregulation of Rab proteins, their regulators, or their effectors has been linked to cancer, rare genetic diseases, and neurodegenerative disorders, underscoring the need to define their molecular mechanisms and disease-relevant roles.^4,5^

Rab GTPases have emerged as key modulators of mitochondrial quality control pathways, particularly in the context of neurodegenerative disease.^6–8^ Mitochondria perform diverse essential functions, and are themselves regulated in part by mitophagy - the selective autophagic turnover of damaged mitochondria - to maintain a healthy mitochondrial pool, to preserve bioenergetics, mitochondrial DNA (mtDNA) integrity, and reactive oxygen species (ROS) balance.^9,10^ The best characterized mitophagy pathway is mediated by serine/threonine PTEN-induced putative kinase 1 (PINK1) and the ubiquitin ligase Parkin, which accumulate on depolarized mitochondria, although several PINK1/Parkin-independent pathways have also been described.^10–12^ Unsurprisingly given Rabs’ central role in vesicular machinery required for autophagic processes, multiple Rab family members, including Rab5, Rab7, Rab8A/B, Rab9, Rab10, and Rab13, have been implicated in canonical and non-canonical mitophagy.^8^ However, a comprehensive, systematic understanding of how individual Rabs regulate mitochondrial quality control is lacking - which is essential both to clarify fundamental mitophagy mechanisms and to reveal targets that could be leveraged to ameliorate mitochondrial dysfunction in neurodegenerative and other Rab-linked diseases.

To address this gap, we performed a comprehensive small interfering RNA (siRNA)-based screen targeting the Rab family, using a HeLa cell line stably expressing the mitophagy reporter mito-mKeima (mt-mKeima) and yellow fluorescent protein-tagged Parkin (YFP-Parkin) to quantify depolarization-induced mitophagy. We identified several novel candidate Rabs that either enhanced or suppressed depolarization-induced mitophagy. We went on to further validate Rab12 as a negative regulator of mitophagy. Given previously reported roles of Rab12 in positive regulation of macroautophagy, we tested whether Rab12 loss reduces baseline, non-selective mitochondrial turnover, thereby allowing accumulation of lower-functioning mitochondria that are more sensitive to mitophagy-inducing stress.^13–18^ Consistent with this model, Rab12 depletion increased mitochondrial content, decreased mitochondrial membrane potential and mitochondrial DNA lesions. These results indicate that one of Rab12’s physiological roles is to restrain mitophagic responses and suggest that its loss lowers the threshold for engagement of the autophagic machinery, pointing to a regulatory function for Rab12 upstream of, or at the level of, autophagosome recruitment/fusion that merits detailed mechanistic follow-up.

## RESULTS

### Family-wide siRNA screen identifies Rab GTPases that modulate mitophagy

We conducted a comprehensive, family-wide siRNA-based screen of Rab GTPases to identify novel regulators of mitophagy. Using an siRNA library targeting sixty-one Rab proteins (**Table 1**), we interrogated mitophagy in a previously described HeLa cell line that stably expresses a mitochondria-targeted mKeima (mt-mKeima).^19^ mt-mKeima is a mitochondria-localized fluorescent protein whose excitation spectrum shifts in a pH-dependent manner when mitochondria are engulfed within acidic lysosomes (pH 4-5), enabling robust image-based quantification of mitophagic flux.^20^ This line was also engineered to express YFP-tagged Parkin, as Parkin is not endogenously expressed by HeLa cells, and localization of YFP-Parkin to mitochondria following mitophagy induction is a secondary validation of active mitophagy.^19^ CRISPR-based screens using this mt-mKeima-expressing HeLa model have previously uncovered core mitophagy regulators and pathways.^21^ Notably, this cell line expresses sufficient endogenous leucine-rich repeat kinase 2 (LRRK2), a kinase that drives phosphorylation of multiple Rabs, to enable detection of changes in endogenous Rabs and expresses most, though not all, Rab family members, under physiological conditions (**Supplementary Table 1; Supplementary Figure 1**).^1^

**Table 1:** siRNA library plate details for Rab family protein targets. siRNA library information for Rab family proteins analyzed in our siRNA mitophagy screen. Plate barcodes for sets of 4 siRNAs against each gene of interest, plate ID, and gene information are listed.

| Plate Barcode | Plate Barcode | Plate Id | Entrez Gene Id | Gene Symbol | Gene Description |
| --- | --- | --- | --- | --- | --- |
| 111146357 | 111146431 | 33_36 | 5861 | RAB1A | RAB1A, member RAS oncogene family |
| 111146389 | 111146463 | 158_161 | 81876 | RAB1B | RAB1B, member RAS oncogene family |
| 111146357 | 111146431 | 33_36 | 5862 | RAB2A | RAB2A, member RAS oncogene family |
| 111146358 | 111146432 | 37_40 | 84932 | RAB2B | RAB2B, member RAS oncogene family |
| 111146354 | 111146428 | 21_24 | 5864 | RAB3A | RAB3A, member RAS oncogene family |
| 111146357 | 111146431 | 33_36 | 5865 | RAB3B | RAB3B, member RAS oncogene family |
| 111146357 | 111146431 | 33_36 | 115827 | RAB3C | RAB3C, member RAS oncogene family |
| 111146357 | 111146431 | 33_36 | 9545 | RAB3D | RAB3D, member RAS oncogene family |
| 111146357 | 111146431 | 33_36 | 5867 | RAB4A | RAB4A, member RAS oncogene family |
| 111146357 | 111146431 | 33_36 | 53916 | RAB4B | RAB4B, member RAS oncogene family |
| 111146357 | 111146431 | 33_36 | 5868 | RAB5A | RAB5A, member RAS oncogene family |
| 111146357 | 111146431 | 33_36 | 5869 | RAB5B | RAB5B, member RAS oncogene family |
| 111146357 | 111146431 | 33_36 | 5878 | RAB5C | RAB5C, member RAS oncogene family |
| 111146357 | 111146431 | 33_36 | 5870 | RAB6A | RAB6A, member RAS oncogene family |
| 111146358 | 111146432 | 37_40 | 51560 | RAB6B | RAB6B, member RAS oncogene family |
| 111146357 | 111146431 | 33_36 | 84084 | RAB6C | RAB6C, member RAS oncogene family |
| 111146357 | 111146431 | 33_36 | 7879 | RAB7A | RAB7A, member RAS oncogene family |
| 111146357 | 111146431 | 33_36 | 338382 | RAB7B | RAB7B, member RAS oncogene family |
| 111146357 | 111146431 | 33_36 | 4218 | RAB8A | RAB8A, member RAS oncogene family |
| 111146357 | 111146431 | 33_36 | 51762 | RAB8B | RAB8B, member RAS oncogene family |
| 111146357 | 111146431 | 33_36 | 9367 | RAB9A | RAB9A, member RAS oncogene family |
| 111146358 | 111146432 | 37_40 | 51209 | RAB9B | RAB9B, member RAS oncogene family |
| 111146357 | 111146431 | 33_36 | 10890 | RAB10 | RAB10, member RAS oncogene family |
| 111146358 | 111146432 | 37_40 | 8766 | RAB11A | RAB11A, member RAS oncogene family |
| 111146357 | 111146431 | 33_36 | 9230 | RAB11B | RAB11B, member RAS oncogene family |
| 111146408 | 111146482 | 231_234 | 201475 | RAB12 | RAB12, member RAS oncogene family |
| 111146357 | 111146431 | 33_36 | 5872 | RAB13 | RAB13, member RAS oncogene family |
| 111146358 | 111146432 | 37_40 | 51552 | RAB14 | RAB14, member RAS oncogene family |
| 111146398 | 111146472 | 194_197 | 376267 | RAB15 | RAB15, member RAS oncogene family |
| 111146357 | 111146431 | 33_36 | 64284 | RAB17 | RAB17, member RAS oncogene family |
| 111146357 | 111146431 | 33_36 | 22931 | RAB18 | RAB18, member RAS oncogene family |
| 111146399 | 111146473 | 198_201 | 401409 | RAB19 | RAB19, member RAS oncogene family |
| 111146357 | 111146431 | 33_36 | 55647 | RAB20 | RAB20, member RAS oncogene family |
| 111146357 | 111146431 | 33_36 | 23011 | RAB21 | RAB21, member RAS oncogene family |
| 111146357 | 111146431 | 33_36 | 57403 | RAB22A | RAB22A, member RAS oncogene family |
| 111146357 | 111146431 | 33_36 | 51715 | RAB23 | RAB23, member RAS oncogene family |
| 111146357 | 111146431 | 33_36 | 53917 | RAB24 | RAB24, member RAS oncogene family |
| 111146361 | 111146435 | 49_52 | 57111 | RAB25 | RAB25, member RAS oncogene family |
| 111146357 | 111146431 | 33_36 | 25837 | RAB26 | RAB26, member RAS oncogene family |
| 111146357 | 111146431 | 33_36 | 5873 | RAB27A | RAB27A, member RAS oncogene family |
| 111146357 | 111146431 | 33_36 | 5874 | RAB27B | RAB27B, member RAS oncogene family |
| 111146355 | 111146429 | 25_28 | 9364 | RAB28 | RAB28, member RAS oncogene family |
| 111146357 | 111146431 | 33_36 | 8934 | RAB7L1<br>(RAB29) | RAB7, member RAS oncogene family-like 1 |
| 111146357 | 111146431 | 33_36 | 27314 | RAB30 | RAB30, member RAS oncogene family |
| 111146357 | 111146431 | 33_36 | 11031 | RAB31 | RAB31, member RAS oncogene family |
| 111146357 | 111146431 | 33_36 | 10981 | RAB32 | RAB32, member RAS oncogene family |
| 111146357 | 111146431 | 33_36 | 9363 | RAB33A | RAB33A, member RAS oncogene family |
| 111146357 | 111146431 | 33_36 | 83452 | RAB33B | RAB33B, member RAS oncogene family |
| 111146357 | 111146431 | 33_36 | 83871 | RAB34 | RAB34, member RAS oncogene family |
| 111146357 | 111146431 | 33_36 | 11021 | RAB35 | RAB35, member RAS oncogene family |
| 111146357 | 111146431 | 33_36 | 9609 | RAB36 | RAB36, member RAS oncogene family |
| 111146357 | 111146431 | 33_36 | 326624 | RAB37 | RAB37, member RAS oncogene family |
| 111146357 | 111146431 | 33_36 | 23682 | RAB38 | RAB38, member RAS oncogene family |
| 111146357 | 111146431 | 33_36 | 54734 | RAB39 | RAB39, member RAS oncogene family |
| 111146357 | 111146431 | 33_36 | 116442 | RAB39B | RAB39B, member RAS oncogene family |
| 111146357 | 111146431 | 33_36 | 142684 | RAB40A | RAB40A, member RAS oncogene family |
| 111146357 | 111146431 | 33_36 | 10966 | RAB40B | RAB40B, member RAS oncogene family |
| 111146357 | 111146431 | 33_36 | 57799 | RAB40C | RAB40C, member RAS oncogene family |
| 111146357 | 111146431 | 33_36 | 347517 | RAB41 | RAB41, member RAS oncogene family |
| 111146392 | 111146466 | 170_173 | 115273 | RAB42 | RAB42, member RAS oncogene family |
| 111146366 | 111146440 | 69_72 | 339122 | RAB43 | RAB43, member RAS oncogene family |

To develop the screen, we assembled an siRNA library targeting the Rab family across fourteen 96-well plates, with each Rab protein targeted by a pool of four distinct siRNAs (**Table 1; Supplementary Table 2**). Each plate included a non-targeting scramble siRNA as a negative control and a PINK1 siRNA as a positive control for impaired mitophagy (**Figure 1A**). After plate siRNA stamping, siRNAs were combined with reagents for reverse transfection of mt-mKeima/YFP-Parkin HeLa cells, followed by a 24-hour (h) incubation to allow target knockdown (**Supplementary Figure 2A, B**). Cells were then treated for 4 h with 30 μM carbonyl cyanide 4-trifluoromethoxy phenylhydrazone (FCCP) or vehicle and imaged live by confocal microscopy to assess mitophagy (**Figure 1A**). Mitophagy was quantified by measuring mt-mKeima puncta that were positive at low pH (4-5) and negative at neutral pH (6-7), corresponding to mitochondria delivered to acidic lysosomes (**Figure 1B, C**). This method of analysis showed that, as expected, FCCP robustly induced mitophagy in scramble-transfected cells (**Figure 1C**).^22,23^ The siRNA designed against PINK1 was used as a functional positive control; PINK1 knockdown was verified through quantitative real-time polymerase chain reaction (qRT-PCR), which showed a >95% decrease in PINK1 gene expression relative to scramble siRNA-transfected control conditions (**Supplementary Figure 2B)**. Loss of PINK1 blocked mitophagy induction, consistent with prior reports (**Figure 1C**).^23,24^

**Figure 1.**
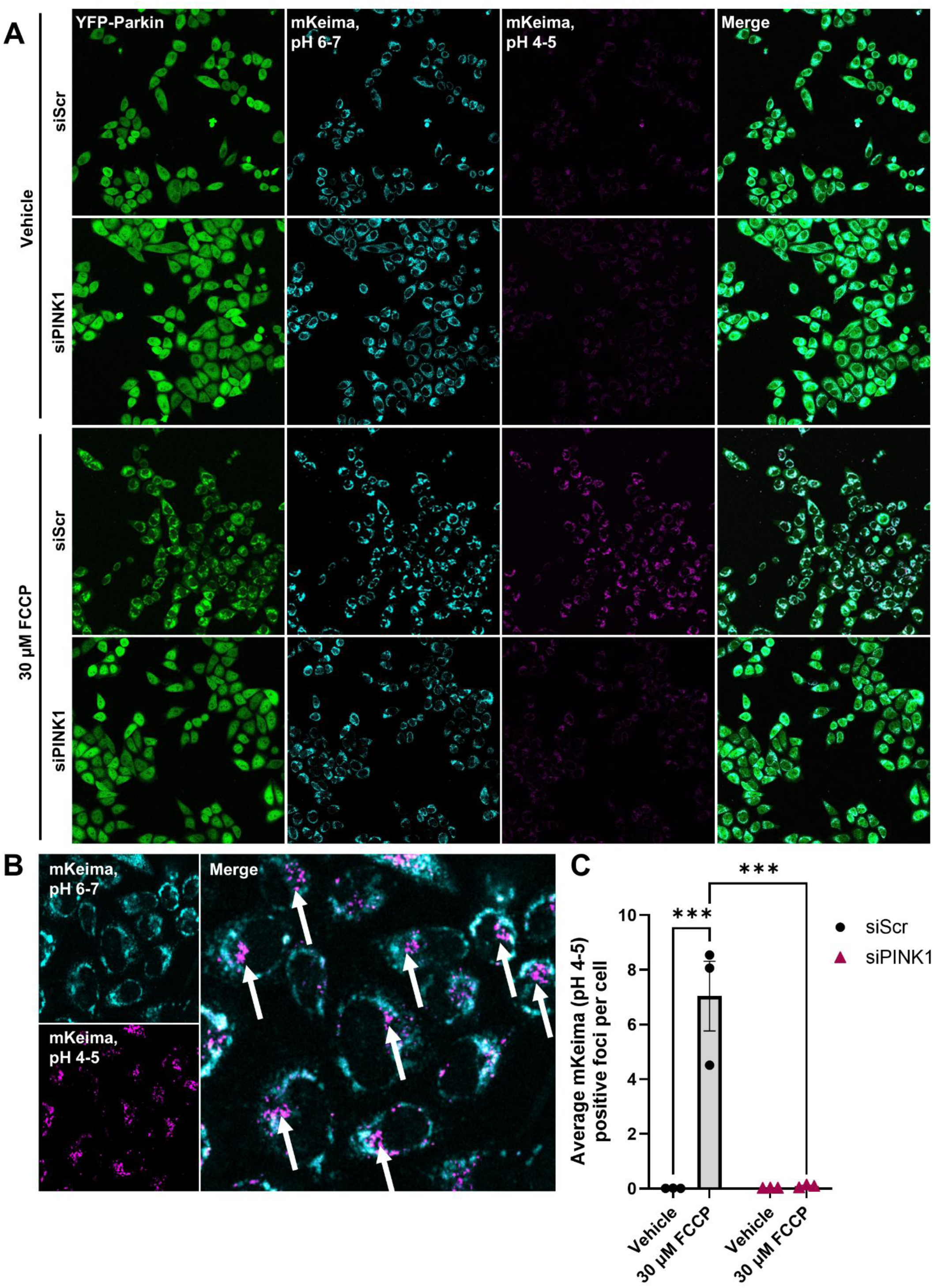
Validation of HeLa cells stably expressing mt-mKeima/YFP-Parkin as a tool for mitophagy analysis. (**A**) Representative 20X confocal images of YFP-Parkin (green), mt-mKeima at neutral pH (cyan), and mt-mKeima at lysosomal pH (magenta) following FCCP-induced mitophagy in mt-mKeima/YFP-Parkin HeLa cells, transfected with a scramble control siRNA (siScr) or an siRNA against PINK1 (siPINK1) for 24 h. (**B**) Mitophagy was analyzed through quantification of puncta positive for lysosomal mt-mKeima (pH 4-5) signal and negative for mt-mKeima at neutral pH (pH 6-7) signal, which are indicated with white arrows. (**C**) This method of analysis was used to quantify mitophagy in mt-mKeima/YFP-Parkin HeLa transfected with siScr or siPINK1 for 24 h, followed by mitophagy induction with a 4 h vehicle control or 30 μM FCCP treatment. \*\*\**p* < 0.005, as determined by two-way ANOVA with Bonferroni’s multiple comparisons test. *n* = 3 biological replicates. Data are presented as mean ± SEM.

After completing validation experiments, we performed the full Rab siRNA-based screen (**Figure 2**), in which mitophagy was quantified as the fold-change in mt-mKeima (pH 4-5)-positive puncta in each Rab siRNA well, normalized to its corresponding non-targeting siRNA scramble plate control. We applied a 50% increase or decrease in fold change as the threshold to nominate Rab GTPases that modulate mitophagy. This 50% cutoff was supported by the observation that siRNAs targeting Rab family members not endogenously expressed in HeLa cells, based on the Human Protein Atlas, did not produce changes within this range (**Supplementary Table 1; Supplementary Figure 3**).^25^ Using this approach, we identified several Rab candidate proteins that impacted mitophagy. Among these, interestingly, only Rab2A knockdown decreased depolarization-induced mitophagy (**Figure 2**). However, knockdown of Rab3C, Rab5A, Rab12, Rab33B, Rab34, Rab36, Rab38, and Rab40A/C increased FCCP-stimulated mitophagy (**Figure 2**).

**Figure 2.**
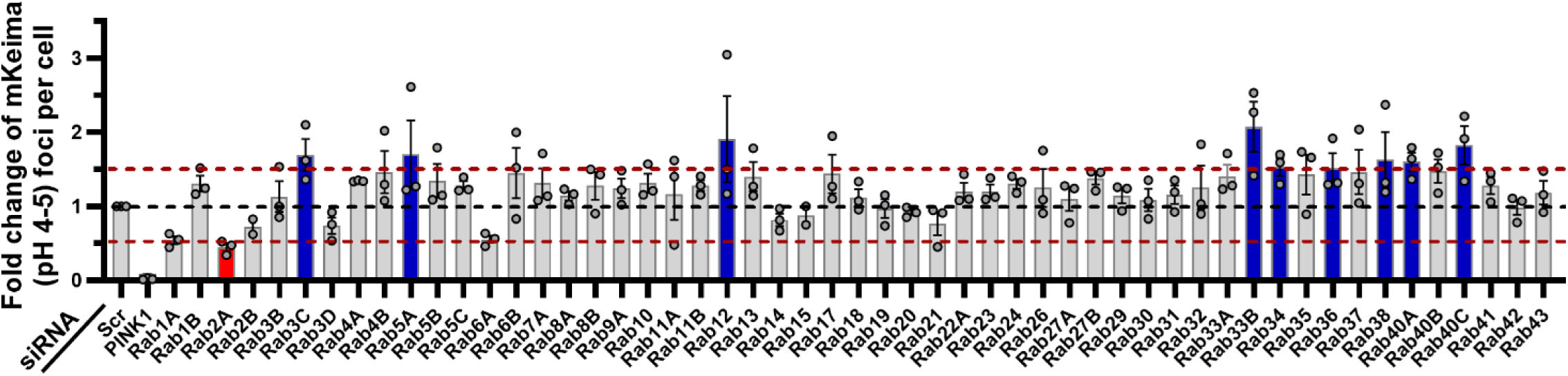
An siRNA-based screen of Rab family proteins for mitophagy effectors. mt-mKeima/YFP-Parkin HeLa cells were reverse transfected with siRNAs against the notated Rab GTPase family members for 24 h. Following transfection, cells were treated with vehicle or 30 μM FCCP for 4h to induce mitophagy. Cells were immediately live-cell imaged on a confocal microscope for mt-mKeima-based mitophagy analysis to analyze impact of Rab target knockdown on mitophagic flux. Results are graphed as average mt-mKeima (pH 4-5) positive foci per cell, normalized to respective plate scramble (Scr) siRNA controls. All plates also included an siRNA against PINK1 as a positive control for mitophagy impairment. Candidates with a ≥50% increase are presented in blue, and those with a ≥50% decrease in mitophagy readout are presented in red. *n =* 3 replicates (except Rab2B and Rab15: *n =* 2 replicates). Data are presented as mean ± SEM.

### Rab12 knockdown enhances basal and depolarization-induced mitophagy

Based on the siRNA-based screen, which identified Rab12 as one of the top candidates for promoting enhanced depolarization-induced mitophagy (**Figure 2**), we selected Rab12 for further analysis. Using an siRNA sequence distinct from those employed in the primary screen, Rab12 knockdown in mt-mKeima/YFP-Parkin HeLa cells reduced Rab12 mRNA levels by ∼60% relative to a non-targeting scramble control, as measured by qRT-PCR (**Figure 3A**). Quantitative immunoblotting demonstrated a corresponding ∼80% decrease in total Rab12 protein in mt-Keima/YFP-Parkin HeLa cells (**Figure 3B, C**). Consistent with the initial findings from the siRNA-based screen (**Figure 2**), Rab12 depletion significantly increased FCCP-induced mitophagy relative to control siRNA-transfected conditions (**Figure 3D, E**). Notably, Rab12 knockdown did not affect cell viability relative to control conditions (**Supplementary Figure 4**). Two-way ANOVA analysis revealed siRab12 and FCCP significantly increased mitophagy compared to controls (**Figure 3E**). In addition, there was a significant interaction effect between siRab12 and FCCP (*p =* 0.0233), supporting that depolarization-induced mitophagy response relative to baseline mitophagy levels was significantly increased with Rab12 knockdown (**Figure 3E**).

**Figure 3.**
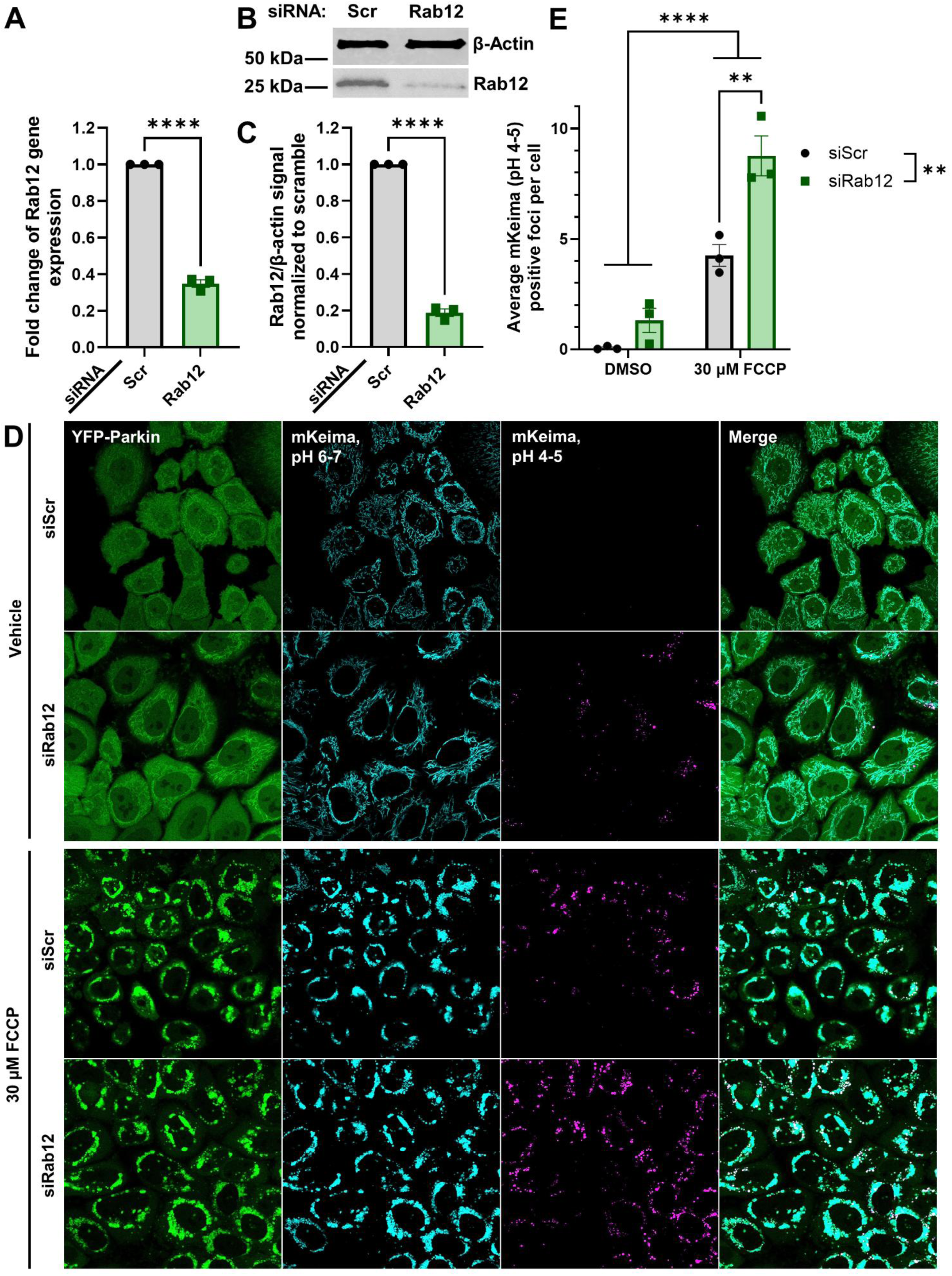
Rab12 knockdown upregulates FCCP-induced mitophagy in mt-mKeima/YFP-Parkin HeLa cells. (**A**) RNA was collected for qRT-PCR analysis of Rab12 mRNA expression in mt-mKeima/YFP-parkin HeLa cells following 72 h transfection with siRNA against Rab12 (siRab12) or the scramble control (siScr). Internal control for normalization was GAPDH. \*\*\*\**p* < 0.0001 determined by unpaired t-test. *n* = 3 biological replicates with three technical replicates each. (**B**) Representative western blot of lysates collected from mt-mKeima/YFP-parkin HeLa cells transfected with siScr or siRab12 assessed for Rab12 levels and β-actin as a loading control. (**C**) Quantification demonstrates Rab12 protein knockdown with Rab12 siRNA by the same protocol. \*\*\*\**p* < 0.0001, as determined by unpaired t-test. *n* = 3 biological replicates. (**D**) Representative 60X confocal images of YFP-Parkin (green), mt-mKeima at neutral pH (cyan), and mt-mKeima at lysosomal pH (magenta) and (**E**) quantification of mt-mKeima mitophagy analysis demonstrated increased levels of mitophagy induced by treatment with 30 μM FCCP with Rab12 knockdown. \*\**p* < 0.01, \*\*\*\**p* < 0.0001, as determined by two-way ANOVA with Bonferroni’s multiple comparisons test. *n* = 3 biological replicates. Data are presented as mean ± SEM.

To explore whether the observed Rab12-dependent increase in depolarization-induced mitophagy is conserved across different cell types, we analyzed basal and depolarization-induced mitophagy in wild-type and Rab12 knockout (Rab12 KO) Raw 264.7 macrophage cell lines.^26^ We employed a pH-sensitive Mitophagy Dye, which increases in fluorescence intensity at lysosomal pH to measure mitophagic flux, and a Lyso Dye to validate lysosomal localization of mitophagy foci. Mitophagy was measured through quantification of punctae with co-localized Mitophagy and Lyso Dye fluorescent signal (mitolysosomes). Two-way ANOVA analysis revealed significant main effects of genotype (wildtype vs. Rab12 KO) and treatment condition. Basal and FCCP-induced mitophagy were significantly increased in Rab12 KO Raw 264.7 cells relative to wild-type control cells to a similar extent (**Figure 4A, B**). Overall, taken together, these findings support a role for Rab12 negatively regulating mitophagy in multiple cell types.

**Figure 4.**
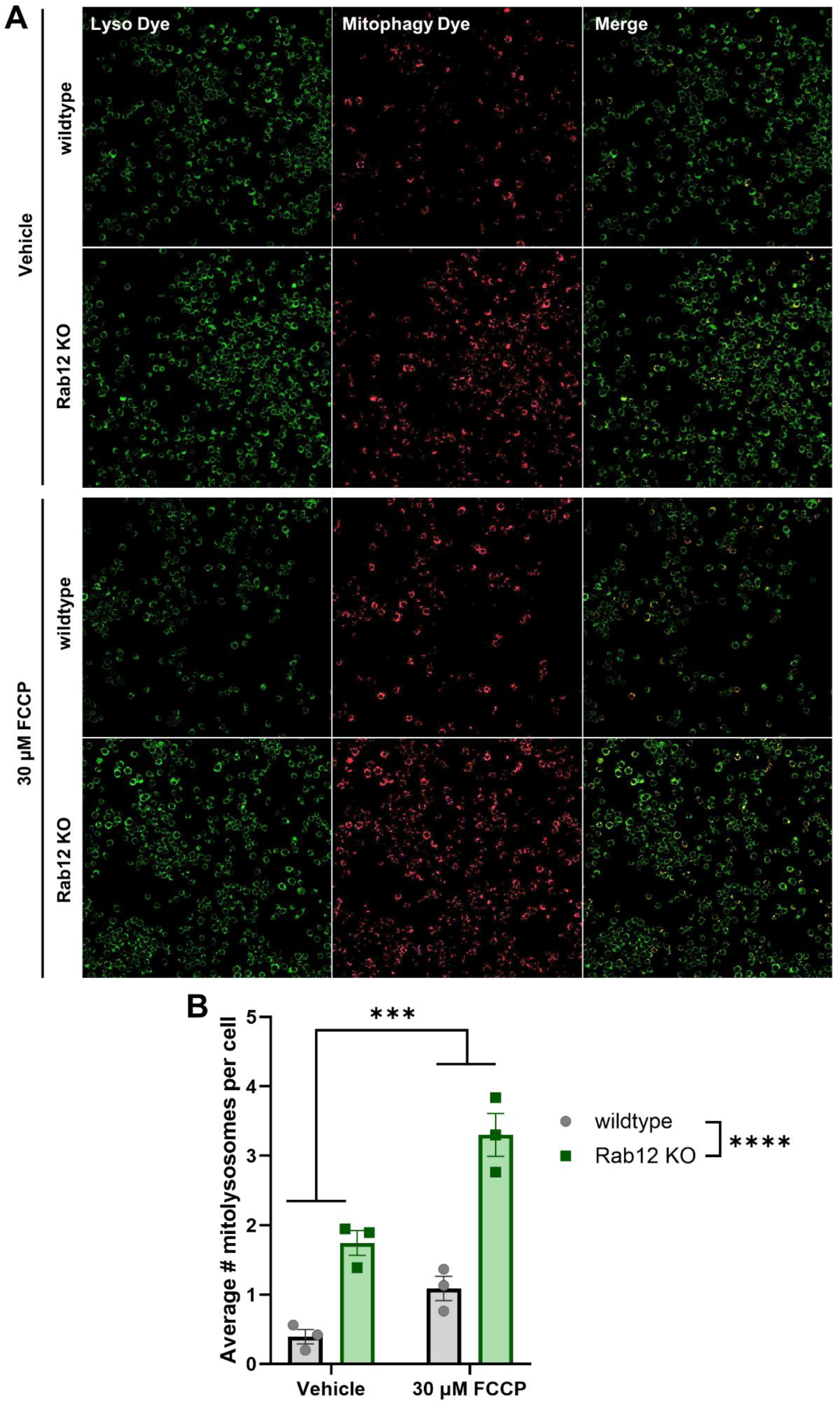
Basal and FCCP-induced mitophagy are upregulated in Rab12 KO Raw 264.7 cells. (**A**) Wildtype and Rab12 KO Raw 264.7 cells were incubated with Mitophagy Dye, and treated with 30 μM FCCP (or vehicle control) for 6 h, treated with Lyso Dye, and imaged to quantify mitophagy. (**B**) Mitophagy was quantified as the average number of mitolysosomes (foci positive for Mitophagy Dye and Lyso Dye signal) per cell. \**\*\*p*<0.001, \*\*\*\**p*<0.0001, as determined by two-way ANOVA. *n* = 3 biological replicates. Data are presented as mean ± SEM.

### Rab12 knockdown impacts mitochondrial membrane potential, mitochondrial content, and mtDNA integrity

Rab12 knockdown has been associated with decreased basal and starvation-induced macroautophagy by multiple groups.^13–18^ We therefore considered that Rab12 knockdown-dependent reduction in macroautophagy may promote the accumulation of excess, poorly functioning mitochondria, leading to enhanced mitophagy following exposure to a mitophagy-specific stressor. To investigate the health of the mitochondrial pool with Rab12 depletion compared to siRNA scramble control conditions, we first employed flow cytometry to measure (1) cellular mitochondrial content, and (2) mitochondrial membrane potential. Neutral pH mt-mKeima fluorescence was used as a readout of mitochondrial content. Interestingly, we found that cells transfected with Rab12 siRNA had significantly increased mitochondrial content compared to those transfected with a control siRNA (**Figure 5A, B**).

**Figure 5.**
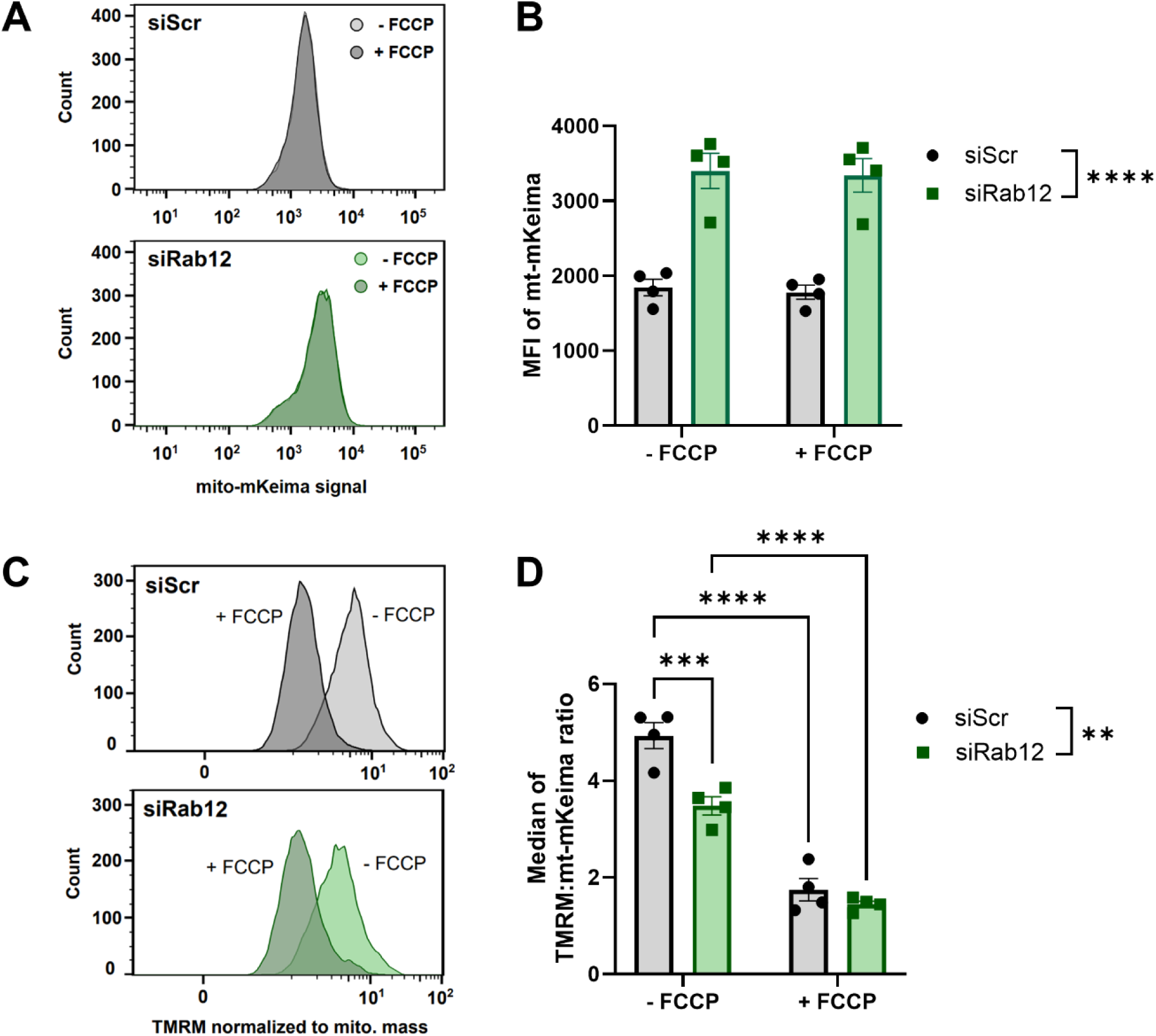
Rab12 knockdown increases mitochondrial content and decreases mitochondrial membrane potential at baseline. (**A** and **B**) mt-mKeima/YFP-Parkin HeLa cells were transfected with siRNA against Rab12 (siRab12) or a scramble siRNA control (siScr) for 72 h. Mitochondrial content was analyzed by quantifying median fluorescence intensity (MFI) of mt-mKeima in siScr- vs siRab12-transfected samples (- FCCP) and both samples were then acutely treated with FCCP and measured once more (+ FCCP). (**C** and **D**) mt-mKeima/YFP-Parkin HeLa cells transfected with siRab12 or siScr for 72 h were collected and stained with TMRM. MFI of TMRM was calculated for siScr and siRab12 samples before and after FCCP treatment to measure mitochondrial membrane potential. \*\**p*<0.01, \*\*\**p*<0.001, *\*\*\*\*p*<0.0001, as determined by two-way ANOVA with Bonferroni’s multiple comparisons test. *n* = 4 biological replicates. Data are presented as mean ± SEM.

Mitochondrial membrane potential was measured using flow cytometry with tetramethyl rhodamine methyl ester perchlorate (TMRM) in mt-mKeima/YFP-Parkin HeLa cells transfected with Rab12 or scramble control siRNA, before and after acute treatment with FCCP. In the absence of exogenous stress, Rab12 siRNA-mediated knockdown significantly decreased mitochondrial membrane potential compared to cells transfected with a scramble siRNA control, when normalized to mitochondrial content, implicating a pool of lower-functioning mitochondria (**Figure 5C, D; Supplementary Figure 5 and 6**). Upon addition of FCCP, both siRNA-mediated scramble control or Rab12 knockdown cells demonstrated significantly decreased mitochondrial membrane potential (**Figure 5C, D**).

**Figure 6.**
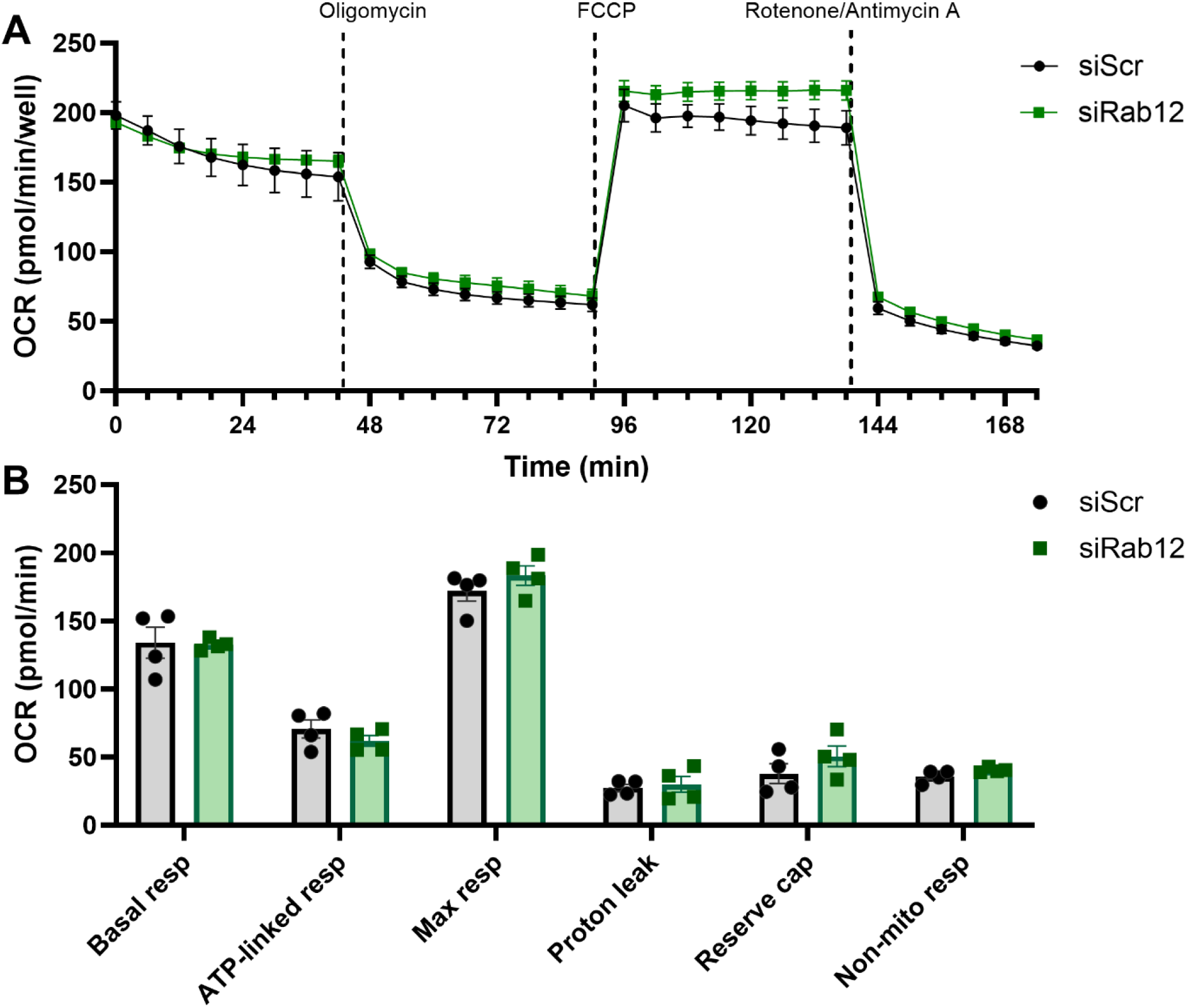
Rab12 knockdown does not alter mitochondrial bioenergetics. (**A**) Representative graph of the Seahorse bioenergetic MitoStress Test (Agilent Biosciences, Santa Clara, CA, USA) conducted on mt-mKeima/YFP-Parkin HeLa cells pre-transfected with an siRNA against Rab12 (siRab12) or a scramble control (siScr) for 72 h prior to Seahorse analysis. (**B**) Extrapolated data from the MitoStress Test of siScr-transfected cells against siRab12-transfected cells, showing basal respiration (basal resp), ATP-linked respiration (ATP-linked resp), maximal respiration (max resp), proton leak, reserve capacity (reserve cap), and non-mitochondrial respiration (non-mito resp). Data was analyzed using a two-way ANOVA with Bonferroni’s multiple comparisons test. *n* = 4 biological replicates. Data are presented as mean ± SEM.

We next determined whether the increased mitochondrial content and accompanied lower mitochondrial membrane potential observed with Rab12 knockdown was associated with changes in bioenergetic function. To do this, we measured oxygen consumption rate (OCR) in mt-mKeima/YFP-Parkin HeLa cells transfected with siRNA to either Rab12 or a scramble control for 72 h, in response to oligomycin (inhibitor of ATP synthase), FCCP (proton ionophore) and rotenone combined with antimycin A (a mitochondrial complex I inhibitor and III inhibitor respectively, **Figure 6A**). Intriguingly, despite an increase in mitochondrial content (**Figure 5A, B**), the basal, ATP-linked or maximal respiration was not different between Rab12 siRNA-mediated knockdown and scramble siRNA conditions in mt-mKeima/YFP-Parkin HeLa cells (**Figure 6A, B**). Reserve capacity, proton leak and non-mitochondrial respiration were also similar between Rab12 knockdown and control cells (**Figure 6A, B**).

Lastly, we analyzed the effect of Rab12 knockdown on mitochondrial genome integrity. To measure mtDNA damage we utilized the Mito DNA_DX_ assay, which has been successfully applied to a variety of cell types and species.^27–35^ We found that mtDNA damage levels were significantly decreased in mt-mKeima/YFP-Parkin HeLa cells with Rab12 siRNA-mediated knockdown relative to the scramble control (**Figure 7A**). Despite this decrease in mtDNA lesion frequency, mtDNA copy number was similar across both conditions (**Figure 7B**). Although the precise mechanisms underlying these findings remain unknown, these results are consistent with our evidence that Rab12 deficiency enhances mitophagy (**Figure 3**). Taken together, Rab12 knockdown results in cells with lower-functioning mitochondria (reduced mitochondrial membrane potential, mitochondrial genome imbalance), which despite a doubling in mitochondrial content do not exhibit increased oxidative capacity.

**Figure 7.**
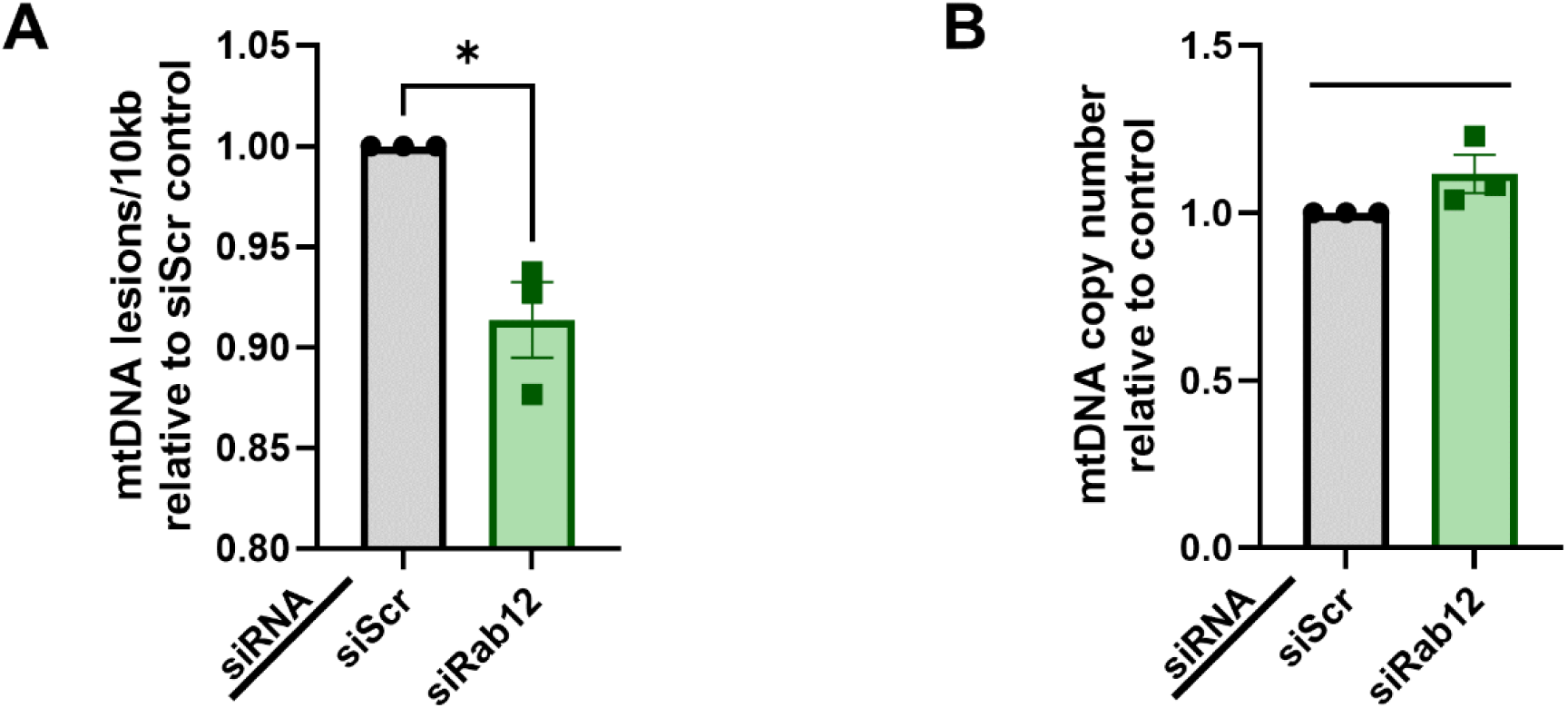
Rab12 knockdown decreases mtDNA damage levels in mt-mKeima/YFP-Parkin HeLa cells. (**A**) mt-mKeima/YFP-Parkin HeLa were transfected with siRNA against Rab12 (siRab12) or a scramble control siRNA (siScr) for 72 h. Mito DNA_DX_ analysis of mtDNA lesion frequency showed that Rab12 knockdown caused a significant decrease in mtDNA lesions relative to siScr transfection. *n* = 3 biological replicates. \**p* < 0.01, as determined by unpaired t-test. (**B**) No differences in steady state of mtDNA copy number was observed between conditions. *n* = 3 biological replicates. Data are presented as mean ± SEM.

## DISCUSSION

Our siRNA-based screen nominated multiple previously unrecognized Rab GTPases as candidate mitophagy effectors. Of the Rabs we identified, only Rab2A, Rab5A, and Rab34 have prior links to mitophagy.^36,37^ Rab2 and Rab34 have been described to be recruited to the mitochondrial membrane following CCCP-induced mitochondrial membrane depolarization and Rab5 has been detected on early endosomes involved in Parkin-dependent mitochondrial clearance.^36,37^ The remaining candidates have established roles across diverse endolysosomal and trafficking processes, including exocytosis and vesicle recycling (Rab3C), endosome-lysosome trafficking and macroautophagy (Rab12), intra-Golgi transport (Rab33B and Rab36), autophagosome maturation (Rab33B), lysosome-related organelle biogenesis (Rab38), and proteasome substrate recognition/binding (Rab40A/C).^14,15,18,38–45^ These functional connections to membrane trafficking and quality-control pathways make them plausible regulators of mitophagy at multiple steps (e.g., organelle recognition, autophagosome formation/maturation, or lysosomal delivery). Future work may validate these candidates in orthogonal assays and delineate the specific stages of mitophagy they influence, including mapping interacting partners, performing epistasis with known mitophagy factors, and testing physiological relevance in disease-relevant cell types or *in vivo* models.

Notably, several Rab proteins previously implicated in mitophagy were not identified in the siRNA-based screen. For example, Rab7A knockdown impaired mitochondrial engulfment by autophagosomes and degradation of mitochondrial markers after valinomycin treatment.^46^ In another study, knockdown of Rab8A impaired mitophagy as measured by COXII depletion after CCCP or oligomycin and antimycin treatment.^37^ Rab10 knockdown reduced mt-mKeima signal and HSPD1 depletion after prolonged valinomycin treatment, while overexpression of FLAG-tagged Rab10 enhanced mitophagy.^47^ Knockdown of Rab9A and Rab9B also suppressed starvation-induced mitophagy in HeLa cells expressing mt-mKeima.^48^ Rab11 has been linked to PINK1/Parkin-dependent mitophagy in *Drosophila melanogaster,* with its knockdown being associated with impairment of mitophagic flux and dopaminergic neuronal function.^49,50^ Rab9 has been reported in alternative mitophagy pathways, with Rab9 knockdown attenuating mitophagy caused by treatment with insulin-like growth factor II in neonatal rat ventricular myocytes, as well as mitophagy induced by the small molecule ALT001 in HEK293 cells.^51,52^ Additional family Rab family members (Rab3A, Rab5, Rab13, Rab14, Rab27, and Rab35) have been linked to canonical or non-canonical mitophagy or related trafficking steps, although knockdown effects on mitophagic flux are not uniformly reported.^36,47,53–59^

Differences in experimental conditions may account for these discrepancies. We used an acute FCCP treatment and mt-mKeima/YFP-Parkin HeLa cells to maximize early, quantifiable mitophagy events while preserving cell viability. Other studies employed more chronic or alternative induction paradigms (longer CCCP/valinomycin exposures, starvation, IGF-II activation, or small-molecule inducers) that can engage alternative mitophagy routes, bulk autophagy, or apoptosis and thus reveal dependencies not evident under acute depolarization. Reporter choice also matters: mt-mKeima reports delivery of mitochondria to acidic compartments and is well suited to high-throughput, endpoint measurement at early timepoints, whereas immunofluorescence or immunoblot detection of mitochondrial marker depletion and assays of autophagosomal engulfment probe different mitophagy stages and kinetics.^20^ Together, these considerations underscore that mitophagy is mechanistically heterogenous and context-dependent. Next steps may include testing newly identified candidates from the current work with varied induction paradigms, extended time courses, and mitophagy reporters to define the specific stages and physiological context in which each Rab operates.

Of the candidates identified in our screen, we prioritized Rab12 for follow-up for several reasons. Recent work from our group found that proteolytically active, lysosome-like structures known as granulovacuolar degeneration bodies in aged PS19 tauopathy mouse brains co-localize with Rab12 phosphorylated at Ser106 (pS106-Rab12) and the mitophagy marker phospho-ubiquitin S65 (pS65-Ub), suggesting a new link between Rab12, mitophagy, and proteinopathy in neurodegeneration.^60^ Although Rab12 has not been previously implicated in selective mitophagy, prior studies report two mechanisms by which Rab12 can positively regulate macroautophagy, either indirectly through regulation of upstream autophagy regulator mTORC1 activity by mediating degradation of the amino-acid transporter PAT4, or directly through facilitation of autophagosome trafficking following ULK-dependent Rab12 activation via Rab12’s guanine exchange factor DENND3.^14,15,18^ First, Rab12 promotes degradation of the amino-acid transporter PAT4, thereby limiting mTORC1 activity; Rab12 knockdown increased phospho-p70S6K and decreased LC3-II levels, consistent with elevated mTORC1 signaling.^15^ Because mTORC1 negatively regulates both macroautophagy and mitophagy, our observation that Rab12 depletion enhances mitophagy is notable in light of this report.^61^ Importantly, the same study found no detectable defect in autophagic flux or lysosomal function by LC3 turnover assays despite changes in autophagy markers.^14^ Second, Rab12 has been proposed to act directly in autophagy by binding LC3^+^ autophagosomes and myosin II to facilitate autophagosome trafficking downstream of ULK1 activation; Rab12 knockdown reduced autophagosome track length and displacement.^14,18^ Together, these findings indicate Rab12 influences multiple aspects of autophagy regulation - both upstream signaling and downstream trafficking - which may help reconcile how Rab12 depletion can paradoxically increase selective mitophagy reported herein, while producing complex effects on macroautophagy in other contexts.

Since Rab12 has been implicated in promoting macroautophagy, our finding that Rab12 knockdown increased mitophagy was surprising and suggested a nuanced regulatory role. Rab12 loss may impair bulk macroautophagic turnover of aged mitochondria, estimated to account for ∼35% of mitochondrial turnover, thereby permitting accumulation of dysfunctional mitochondria under basal conditions.^62^ This model is supported by our data showing increased baseline mitochondrial content and reduced mitochondrial membrane potential after Rab12 depletion. However, one limitation for mt-mKeima signal intensity measured by flow cytometry is the inability to differentiate whether increased mitochondrial content reflects elevated number of mitochondria and/or increased mitochondrial mass from mitochondrial swelling. It is also possible that the observed increase in mitochondrial content could be attributable to an accumulation of mitochondria within autophagosomal and/or lysosomal compartments. Rab12 knockdown has been reported to impair both autophagosome trafficking and autolysosome formation, which may lead to the accumulation of mitochondria-autophagosomes in which mt-mKeima may be detectable, and observed increases in mitophagy could be primarily driven through the engagement of non-canonical mitochondrial clearance (e.g., mitochondrial-derived vesicles [MDVs] or endosome-mediated mitophagy) that bypass defects in autophagosome trafficking.^13,14^ Accumulation of mitolysosomes may also drive an increase in observed mitochondrial content. While prior reports suggest that Rab12 knockdown does not ablate lysosomal function under basal or starvation-induced autophagy conditions, even small lysosomal changes, including minute lysosomal pH changes within the pH 4-5 range, have the propensity to greatly impact protein half-life and catabolism, and this could also promote accumulation of mitochondrial signal through slowed turnover in spite of lysosomal delivery as observed through mitophagy analysis.^15,63,64^

An additional possible mechanism for Rab12-dependent mitochondrial mass accumulation involves increased biogenesis. Mitochondrial depolarization and oxidative stress caused by cytochrome c oxidase deficiency were shown to drive mitochondrial biogenesis in rat fibroblasts, causing an increase in detected mitochondrial content and similar OCR levels relative to control conditions.^65^ It is plausible that Rab12 depletion-dependent mitochondrial dysfunction is driving biogenesis in a similar manner, leading to increased mitochondrial content and functional respiratory compensation, as supported by indistinguishable OCR levels between control and Rab12 knockdown conditions. Further, the increase in mitochondrial mass was not accompanied by an increase in mtDNA content. Rather we observed a decrease in mtDNA damage levels, which is an integrated readout of ROS, mtDNA replication/repair and mitophagy.^66,67^ Collectively, our findings potentially point to an increased turnover or purge of mtDNA via mitophagy, however future analysis may distinguish whole cell changes from mtDNA copies per mitochondria, which is critical given that depletion of mtDNA copy number per mitochondria can act as both a consequence and a driver of mitochondrial dysfunction.^68,69^ Mitochondrial content has been associated with a linear increase in bioenergetic capacity, but a corresponding increase in OCR was not observed under conditions of Rab12 depletion which enhanced mitochondrial content, supporting the existence of a larger, lower-functioning mitochondrial pool and/or an increase in mitochondrial content to energetically compensate for mitochondrial dysfunction.^70–72^

Upon application of a mitophagy-specific stressor, the accumulated pool of lower-functioning mitochondria would be expected to drive greater mitophagic flux, consistent with the increased mt-mKeima delivery to acidic compartments we observed. As noted in the literature discussed above, Rab12 knockdown attenuated but did not ablate basal or induced autophagy, providing a mechanistic rationale for why we still observed a net increase in mitophagic flux, which is generally dependent upon the function of autophagic machinery (autophagosome trafficking/fusion with lysosomes, lysosomal function), even with Rab12 knockdown.^15^ Future work may dissect autophagosomal and lysosomal responses to general versus mitophagy-specific stress in Rab12-deficient cells and employ MDV and endocytic markers to clarify the mechanisms of mitochondrial engulfment and lysosomal delivery.

Mutations in *LRRK2* are the most common cause of autosomal dominant familial Parkinson’s disease (PD).^73,74^ The discovery that multiple Rab GTPases are LRRK2 substrates has opened important avenues for studying PD-relevant pathways.^75^ Among LRRK2’s Rab substrates, Rab12 (like Rab29) is both a substrate and an upstream regulator of LRRK2 kinase activity, and recent work highlights functional interplay between Rab12 and LRRK2.^76–78^ Rab12 localizes to lysosomal membranes following lysosomal stress or in cells expressing PD-linked *LRRK2* variants, facilitating LRRK2 recruitment and enhancing LRRK2-dependent phosphorylation of Rab10, consistent with a role for Rab12 in amplifying LRRK2 activation by pathogenic variants.^77^ Rab12 phosphorylation and its interaction with RILPL1 were found to be key mediators in perinuclear lysosomal clustering driven by *LRRK2-Y1699C,* linking Rab12 to lysosomal transport defects.^79^ Beyond lysosomal effects, Rab12 has been reported to modulate centrosome homeostasis and primary ciliogenesis in

LRRK2-G2019S models, acting as a negative regulator of ciliogenesis upon its interaction with LRRK2, and Rab12 deletion rescued the centrosome/cilia defects.^78^ The convergence of Rab12’s roles in LRRK2-driven lysosomal, centrosomal, and ciliogenesis defects with our demonstration that Rab12 constrains mitophagy raises the compelling possibility that Rab12 is a critical nexus linking LRRK2 pathology to both autophagic and mitochondrial dysfunction across multiple PD-linked *LRRK2* variants. Taken together, our study identifies multiple mitophagy regulators and establishes Rab12 as a conserved negative regulator whose loss promotes accumulation of lower-functioning mitochondria and sensitizes cells to mitophagy-inducing stress, linking Rab-dependent trafficking to mitochondrial quality control with implications for neurodegenerative disease.

## MATERIALS AND METHODS

### Mammalian cell culture, sorting and treatments

The mt-mKeima/YFP-Parkin HeLa cell line was obtained from Dr. Richard Youle, generated via retroviral and lentiviral transfection, as described previously.^19^ Due to heterogeneous levels of mt-mKeima and YFP-Parkin fluorescence, cells were sorted and enriched using a Cellenion cellenONE^®^ image-based single cell sorter. Clones with similar fluorescence levels of both proteins were selected for expansion and use in experiments, with separate sorting clones used for siRNA screening and target validation (**Supplementary Figure 7**). The mt-mKeima/YFP-Parkin HeLa cells were maintained in Dulbecco’s Modified Eagle’s Medium (DMEM) with high glucose, no glutamine (Thermo Fisher Scientific, 11960044) supplemented with 10% Fetal Bovine Serum (VWR, 97068-085), 1% 4-(2-hydroxyethyl)-1-piperazineethanesulfonic acid (HEPES; Thermo Fisher Scientific, 15630080), 1% sodium pyruvate (Thermo Fisher Scientific, 11360070), 1% Minimum Essential Medium - Non-Essential Amino Acids (Thermo Fisher Scientific, 11-140-050), 1% Gibco GlutaMax (Thermo Fisher Scientific, 35050061), 1% Gibco Antibiotic-Antimycotic (Thermo Fisher Scientific, 15240062), in an incubator at 37°C with 5% CO_2_. Wildtype and Rab12 KO Raw 264.7 macrophages were generated via CRISPR/Cas9 as described previously.^26^ Raw264.7 cells were maintained in DMEM with high glucose, pyruvate (Thermo Fisher Scientific, 11995065) supplemented with 10% Fetal Bovine Serum, 1% penicillin/streptomycin, in an incubator at 37°C with 5% CO_2_.

For siRNA transfection experiments, cells were cultured in Antibiotic-Antimycotic-free medium from plating until time of collection. Transfections were performed using Lipofectamine RNAiMAX transfection reagent (Fisher Scientific 13-778-150) diluted in OptiMEM reduced serum medium (Thermo Scientific 31985070) as described below. For experiments involving mitophagy induction, we treated cultures with 30 μM FCCP (Abcam, ab120081) or a v/v DMSO vehicle control (Sigma-Aldrich, D2650) in complete medium for 4 h prior to imaging.

### Rab GTPase siRNA screen

siRNAs against Rab family proteins were pooled from the Qiagen Whole Genome siRNA Set V1.0. The Qiagen Whole Genome library is provided as pre-set 384-well plate arrangements, so we selected individual 96-well quadrants within plates from the library containing siRNAs against Rab family members for use in this screen. Fourteen 96-well plate layouts (barcodes denoted in **Table 1**) were utilized in triplicate for screening, with each plate containing 1 to 43 Rab family member siRNAs and each well containing a pool of 4 siRNAs targeting a single gene. Within the siRNA library utilized, 4 of the 66 identified Rab family members were not included -Rab1C, Rab6D, Rab44, and Rab45. Rab46 was included in the siRNA library but was omitted from this screen due to its lack of expression in HeLa and was not included as a negative control in **Supplementary Figure 3** due to not being grouped on a plate with other Rab protein candidates. siRNA was resuspended in 10 μL UltraPure™ water (Invitrogen, 10977015) to a final 200 nM siRNA concentration. 96-well plates with a glass-like polymer coverslip bottom (Cellvis, P96-1.5P) were stamped using the Bravo Automated Liquid Handling Platform (Agilent Technologies) with 2 pmol Qiagen library siRNA containing the listed members of the Rab GTPase family of proteins (**Table 1**). 2 pmol control siRNAs - scramble siRNA (Qiagen, 1027280) and siPINK1 (Qiagen, 1027417, Geneglobe ID SI03104878) were added to each experimental plate in 10 μL water (200 nM siRNA concentration). Technical triplicates of each plate for both FCCP treated and DMSO conditions were used for this siRNA screening.

A working mix of Lipofectamine RNAiMAX transfection reagent (Fisher Scientific 13-778-150) and Opti-MEM reduced serum medium (Thermo Scientific 31985070) was prepared by diluting 300 μL RNAiMAX in 9.7 mL OptiMEM. A multichannel pipette was used to manually add 10 μL of the RNAiMAX-OptiMEM mix to each well, for a final volume of 0.3 μL RNAiMAX per well. Plates were centrifuged, followed by a 20-minute incubation at room temperature. Following incubation, mt-mKeima/YFP-Parkin HeLa (3 to 6 passages from thaw) were plated using a Matrix® WellMate® (Thermo Scientific) at 7000 cells in 80 μL media per well, for a final well volume of 100 μL and siRNA concentration of 20 nM. Cells were incubated at 37°C, 5% CO_2_ for 24 h. Transfected cells were then treated with 30 μM FCCP (Abcam, ab120081) or DMSO vehicle control (Sigma-Aldrich, D2650) for 4 h. One to two biological replicates were utilized for each condition for each plate. Technical triplicates of one or two identical plates and conditions derived from one or two biological replicates were plated, treated, and imaged simultaneously.

### Mitophagy imaging and analysis

Individual wells from plates were imaged through live-cell imaging with 37°C, 5% CO_2_ incubation using a Zeiss 880 confocal workstation (ZEISS) with a 20x objective and Argon/2 488 nm and 561 nm Diode lasers. Only wells containing siRNAs against Rab targets (plus plate scramble siRNA and siPINK1 controls) were imaged. To expedite the live imaging process and reduce time between imaging of technical triplicates, stage and plate positioning features in Zen Black edition were used to automatically image wells on each plate. A single plane was taken from the center of each well of interest for analysis.

Images were analyzed using QuPath Open Software for Bioimage Analysis v.0.5.1. Raw files from Zen Black edition microscopy software obtained during confocal image capture were used as input, and thresholding values for detection of puncta positive for staining signal were optimized using the Pixel Classification feature in QuPath software, with pixels of intensity value above the identified background threshold classified as positive detections for further intensity and area quantification. Intensity and shape features were then calculated in QuPath, exported to Microsoft Excel for further analysis, and used to quantify positive mt-mKeima (pH 4-5) puncta, which was normalized to cell count per image to obtain a value of positive mt-mKeima (pH 4-5) puncta per cell. Values for each target were normalized to their respective plate scramble siRNA control to account for plate-to-plate variability. A threshold of a 50% increase or decrease in puncta per cell was designated to identify targets predicted to be involved in regulation (negative or positive, respectively) of mitophagy.

### siRNA screen Rab12 validation

To validate effects of Rab12 knockdown on mitophagy, we optimized a separate transfection protocol to maximize Rab12 knockdown in a 6-well format. mt-mKeima/YFP-Parkin HeLa were plated in a 6-well plate for cell pellet collection or a 35-mm polystyrene glass bottom dish for imaging at a density of 40,000 cells per well. Plated cells were grown at 37°C, 5% CO_2_ for 24 h and were then transfected with 20 nM Rab12 siRNA (Qiagen, 1027417, GeneGlobe ID SI05113171) or a scramble control (Qiagen, 1027280) using OptiMEM and RNAiMAX (6 μL per well). Media was changed to complete medium after 24 h of transfection, followed by an additional 48 h of growth for a total of 72 h of transfection prior to collection. At the time of harvest, plates used for imaging were treated with 30 μM FCCP or a DMSO vehicle control for 4 h, followed by live-cell imaging with 37°C, 5% CO_2_ incubation on a Zeiss 880 inverted confocal workstation with a 63x (1.4NA) oil immersion objective and Argon/2 488 nm and 561 nm Diode lasers. Mitophagy was analyzed as described above using QuPath software. For knockdown validation, cell pellets and lysates were harvested from 6-well plates and processed as described below.

### RNA extraction, cDNA conversion, and real-time quantitative PCR

For mt-Keima/YFP-Parkin HeLa cells, two 6-well plate wells were combined for each treatment analysis. Cells were trypsinized and centrifuged for 5 minutes at 200 x g. Cell pellets were stored at -80°C. Cell pellets were collected from three separate biological replicates for the data presented. RNA was extracted using the Qiagen RNeasy Mini Kit (Qiagen, 74104) according to manufacturer protocols and RNA samples were quantified using a NanoDrop 2000c Spectrophotometer (Thermo Fisher Scientific). 1 μg of total RNA were used for cDNA synthesis using the Qiagen RT^2^ First Strand Kit (Qiagen, 330404). cDNA was diluted 1:10 in autoclaved nuclease-free water (Qiagen, 129114) to obtain a final concentration of 5 ng/μL. Quantitative Real-Time PCR (qRT-PCR) was performed using the Applied Biosystems™ PowerUp™ SYBR™ Green Master Mix (Thermo Fisher Scientific, A25742) and validated primers (Qiagen, 330001) for human PINK1 (GeneGlobe ID PPH20890B-200), human Rab12 (GeneGlobe ID PPH08842A-200), and human GAPDH as a housekeeping gene (GeneGlobe ID PPH00150F-200). Each primer used was diluted 1:1 in nuclease-free water for use in the Mastermix. Mastermixes for each primer were made using 10 μL SYBR™ Green Master Mix per reaction, 1 μL per reaction of 1:1 Qiagen primer, and 7 μL per reaction of nuclease-free water, for a final volume of 18 μL per reaction. Three reactions were made for each primer for each sample and loaded into a MicroAmp Optical 96-well plate with barcode (Applied Biosystems, 4306737), and 2 μL of cDNA was added to each reaction for a final reaction volume of 20 μL. The microplate was sealed with MicroAmp Optical Adhesive Film (Thermo Fisher Scientific, 4311971), vortexed for 1 minute, centrifuged for 20 seconds, and was run on a QuantStudio™ 3 PCR System (A28132). For qRT-PCR parameters, we utilized parameters involving a hold step (50°C for 2 minutes, 95°C for 2 minutes), a PCR cycle (40 cycles of 95°C for 15 seconds, 60°C for 2 minutes), and a final melt curve (95°C for 15 seconds, 60°C for 1 minute, 95°C for 1 second). Each sample was run in triplicate. Data was analyzed using Thermo Fisher Scientific’s Thermo Fisher Connect platform to obtain Cq values, and Microsoft Excel was used to calculate ΔCt, ΔΔCt, and fold change of expression of genes of interest relative to the housekeeping gene of the control sample.

### Western immunoblot analysis and antibodies

For measurement of LRRK2 protein levels, mt-mKeima/YFP-Parkin HeLa were plated at 250,000 cells per well in two 6-well plate wells and lysates were harvested following 24 h of growth. For Rab12 knockdown validation, two 6-well wells per condition of mt-mKeima/YFP-Parkin HeLa cells were plated at 40,000 cells per well. Cells were transfected with Rab12 or scramble control siRNA as described, with cell lysis and protein collection occurring after a total of 72 h of transfection. At the time of harvest, ∼2 million cells were collected and resuspended in 75 μl lysis buffer [RIPA buffer (Sigma-Aldrich, R0278-50ML) with 1% Halt phosphatase inhibitor cocktail (Thermo Fisher, 78420) and 5% protease inhibitor cocktail (Sigma-Aldrich, P8340) by volume]. Lysates were incubated on ice for 10 minutes, followed by centrifugation at 4°C, 10,000 x g for 15 minutes to precipitate cell debris, and supernatants were collected for use in Western immunoblot analysis. 150 μg of protein were loaded for LRRK2 blots, and 100 μg were loaded for Rab12 blots. Protein samples were boiled at 100°C for 5 minutes with dithiothreitol (Bio-Rad, 1610610) and NuPage LDS Sample Buffer (Thermo Fisher Scientific, NP0007) prior to loading into a 4-20% Mini-PROTEAN TGX Stain-Free precast gels (Bio-Rad, 4568096) and separation by SDS-PAGE. Separated proteins were transferred onto a nitrocellulose membrane (Bio-Rad, 1704271) and blocked in 5% w/v nonfat dry milk in 1x PBST (0.5% Tween 20) for 30 minutes. For our analysis, we utilized the following antibodies: rabbit anti-LRRK2 (MJFF2 c41-2), (Abcam, ab133474, 1:2000), sheep anti-Rab12 (MRC-PPU, SA227, 0.5 μg/mL), and mouse β-actin (Novus Biologicals, NBP1-47423, 1:10,000). Membranes for LRRK2 or β-actin measurement were probed with IRDye donkey anti-mouse and anti-rabbit fluorescent-labeled secondary antibodies (LI-COR, 926-32212, 926-68072) in 5% milk TBST for 45 minutes and scanned using an Odyssey Imaging scanner (LI-COR). Membranes for Rab12 measurement were probed with AffiniPure® Donkey Anti-Sheep IgG (H+L) (Jackson ImmunoResearch, 713-005-003) in 5% milk TBST for 45 minutes, followed by development with Crescendo ECL reagent (Millipore Sigma, WBLUR0100) and imaging by Chemidoc MP platform (Bio-Rad). Fluorescent blots were quantified using ImageStudio (Adobe) and ECL blots were quantified using ImageLab (Bio-Rad). Rab12 levels were normalized to β-actin loading controls for each sample. For knockdown verification, samples from three separate biological replicates were analyzed to determine mean knockdown of protein levels following transfection with Rab12 or scramble siRNA.

### Mitophagy analysis in wildtype and Rab12 KO Raw264.7 cells

Wildtype or Rab12 KO Raw264.7 cells were plated on 35-mm polystyrene glass bottom dishes for imaging at a density of 750,000 cells per well, followed by a 24 h incubation prior to mitophagy analysis. Mitophagy was analyzed using Dojindo’s Mitophagy Detection Kit (MD-01) according to the manufacturer protocol. Cells were washed two times with serum-free medium prior to a 30 min incubation with Mtphagy Dye (Dojindo, MD-01) in serum-free medium. Complete medium was used to prepare a 30 µM FCCP or DMSO control treatment solution, which was used to treat cells for 6 h. Following mitophagy induction, cells were washed two additional times with serum-free medium, and a working solution of Lyso dye (Dojindo, MD-01) was prepared in serum-free medium and used to treat cells for 30 min. Following Lyso Dye incubation, cells were washed two times with HEPES (Thermo Fisher Scientific, 15630080), followed by adding fresh HEPES to culture dishes for imaging. Cells were imaged with 37°C, 5% CO_2_ incubation on a Zeiss 880 inverted confocal workstation with a 20x objective and Argon/2 488 nm and 561 nm Diode lasers.

Images were analyzed using QuPath Open Software for Bioimage Analysis v.0.5.1. Raw files from Zen Black edition microscopy software obtained during confocal image capture were used as input, and thresholding values for detection of puncta positive for Mtphagy Dye signal were optimized using the Pixel Classification feature in QuPath software, with pixels of intensity value above the identified background threshold classified as positive detections for further intensity and area quantification. Intensity and shape features were then calculated in QuPath, exported to Microsoft Excel for further analysis, and used to quantify puncta positive for Mtphagy Dye and Lyso Dye signal, which was normalized to cell count per image to obtain a mean value of mitolysosomes per cell.

### Flow cytometry for measuring mitochondrial membrane potential

mt-mKeima/YFP-Parkin HeLa cells were plated and transfected with either a scramble or Rab12-targeting siRNA as described above. Seventy-two h after transfection, cells were harvested by trypsinization, neutralized with complete media, and stained for flow cytometry. Cells were aliquoted (2.0 - 4.0 x 10^5^ per sample) and washed once with PBS, pH 7.4 (Thermo Fisher Scientific, 10010049), and resuspended at a concentration of 4.0 x 10^5^ cells/mL in FACS buffer (1X PBS [Fisher Scientific, BP3994] with 1% BSA [Thermo Scientific, A13100]) with a final concentration of 15 nM TMRM (Invitrogen, T668) and 1.0 ug/mL DAPI (Sigma, D9542) for 15 minutes protected from light. Both dyes were left in cell suspension during flow acquisition to maintain steady-state TMRM equilibration. Samples were acquired on a BD LSRFortessa X-20 flow cytometer. TMRM was excited with a 561 nm laser and detected using a 585/15 bandpass filter. The neutral form of mt-mKeima was excited with a 488 nm laser and detected using a 530/30 bandpass filter. DAPI was excited with a 405 nm laser and detected using a 610/20 bandpass filter. Events were gated to exclude debris, and 10,000 events were acquired from this population. After acquisition, each tube was spiked to a final concentration of 30 uM FCCP or vehicle, incubated for 5 min, and re-acquired without washing.

In FlowJo, compensation matrices were generated for each experiment using a TMRM single-stained control tube of wildtype HeLa cells; a single-stained tube for DAPI containing a 1:1 mixture of live and heat-killed cells (65°C for 5 min followed by 1 min on ice); and an unstained tube of mt-mKeima/YFP-Parkin HeLa cells. An unstained tube of wildtype HeLa cells was used as the negative population for TMRM and mt-mKeima. Cells were gated in FlowJo by forward and side scatter to exclude debris, followed by singlet discrimination using FSC-A versus FSC-H. Live cells were defined by excluding DAPI-positive events. For the live, singlet population, the fluorescence intensity for TMRM was normalized to the fluorescence intensity for mt-mKeima to control for mitochondrial mass. The median normalized TMRM value for each sample was calculated from the resulting ratio distribution.

### Measurement of cellular oxygen consumption rate (OCR)

mt-mKeima/YFP-Parkin HeLa cells were plated at 3,000 cells per well in Agilent Seahorse XFe96/XF Pro microplates (Agilent Technologies, 103792-100), transfected with Rab12 or scramble control siRNA as described, and allowed to grow for 72 h prior to the respiration measurement. On the day of the Seahorse assay, growth media was removed and replaced with Seahorse XF DMEM assay medium (Agilent Technologies, 103575-100) supplemented with 10 mM glucose, 1 mM pyruvate, and 2 mM L-glutamine, then incubated for 1 h in a 37°C non-CO_2_ incubator.

OCR was then measured using the Seahorse XFe96 Extracellular Flux Analyzer (Agilent Technologies) at 6 minute increments starting with baseline, followed by sequential injections of three electron transport chain (ETC) inhibitors, oligomycin, FCCP, and rotenone/antimycin A. 25µL of each ETC inhibitor was added to the appropriate port of the Seahorse cartridge. Port A contained 10µM oligomycin (Millipore Sigma, O4876) for a final in-well concentration of 1.25µM, 0.1% DMSO. Port B was loaded with 7.2 µM FCCP (Cayman Chemical, 15218), for a final in-well concentration of 0.8 µM, 0.1% DMSO. Port C was loaded with 15 µM Rotenone/Antimycin A, (Rotenone Millipore Sigma, R8875, Antimycin A, Millipore Sigma, A8674) for a final in-well concentration of 1.5 µM each, 0.1% DMSO. 26-28 technical replicates were collected for each condition per Seahorse plate. OCR measurements allowed quantification of bioenergetic parameters, including basal respiration, ATP-linked respiration, maximal respiration, proton leak, reserve respiratory capacity, and non-mitochondrial respiration.

### DNA isolation, quantitation, and mitochondrial DNA damage analysis

For mtDNA damage analysis, we plated three 6-well wells per condition at 40,000 mt-mKeima/YFP-Parkin HeLa cells per well, transfected cells at 24 h with Rab12 or scramble control siRNA as described, and allowed cells to grow for 72 h post-transfection prior to DNA collection. We harvested cells (three wells per condition combined for DNA isolation), isolated and quantified genomic DNA, and performed the Mito DNA_DX_ assay for mitochondrial genome integrity and mtDNA copy number as previously described.^27^ mtDNA damage analysis is derived from three biological replicates performed in technical triplicate.

### Statistical Analyses

Data were analyzed using GraphPad Prism software (version 10.6.0). Mitophagy, mitochondrial membrane potential, mitochondrial content, and OCR were analyzed via ordinary two-way ANOVA with Bonferroni’s post hoc multiple comparisons test, with significance defined by *p* value ≤ 0.05. qRT-PCR fold change of Rab12 gene expression, Rab12 protein expression, mtDNA damage levels and mtDNA copy number were analyzed using unpaired t-test, with significance defined by *p* value ≤ 0.05. For all graphs, values and error bars are presented as mean ± standard error of the mean (SEM).

## RESOURCE AVAILABILITY

### Lead contact

Requests for further information and resources should be directed to and will be fulfilled by the lead contact, Laurie H. Sanders.

### Materials availability

This study did not generate new unique reagents.

### Data and code availability

#### Data

All data reported in this paper will be shared by the lead contact upon request.

#### Code

This paper does not report original code.

#### Additional information

Any additional information required to reanalyze the data reported in this paper is available from the lead contact upon request.

## ACKNOWLEDGEMENTS

The mt-mKeima/YFP-Parkin HeLa cell line used in these studies were an appreciated gift from the laboratory of Richard Youle, Ph.D. at the National Institute of Neurological Disorders and Stroke. We would like to thank Matt Scaglione, Ph.D. and members of his lab for assistance with sorting of mt-mKeima/YFP-Parkin HeLa cell lines prior to screening. We also thank the staff of the Duke University Microscopy Core, the Duke University Functional Genomics Core, and the Duke Cancer Institute Flow Cytometry Shared Resource for technical assistance in accomplishing this work. This work was supported by National Institutes of Health (NIH) R35ES035049 and T32ES021432 to J.N.M., and R01NS119528 to L.H.S.

## AUTHOR CONTRIBUTIONS

**T.R.:** Conceptualization, formal analysis, investigation, visualization, writing - original draft preparation, writing - review & editing; **A.G.:** Investigation, formal analysis, visualization, writing - review & editing; **A.B.:** Investigation, writing - review & editing; **I.T.R.:** Investigation, formal analysis, writing - review & editing; **C.F.:** Investigation, writing - review & editing; **T.M.:** Investigation, resources, writing - review & editing; **J.L.:** Investigation, writing - review & editing; **S.A.B.:** Investigation, writing - review & editing; **I.B.:** Investigation, writing - review & editing; **S.Y.K.:** Investigation, resources, writing - review & editing; **X.N.:** Investigation, formal analysis, writing - review & editing; **A.B.W.:** Conceptualization, resources, writing - review & editing; **J.N.M.:** Conceptualization, funding acquisition, resources, supervision, writing - original draft preparation, writing - review & editing; **L.H.S.:** Conceptualization, funding acquisition, supervision, writing - original draft preparation, writing - review & editing.

## SUPPLEMENTARY MATERIAL

**Supplementary Table 1:** Endogenous mRNA expression levels of Rab family protein in HeLa. Gene RNA expression levels, presented as normalized transcripts per million (nPTM), of each Rab GTPase family member analyzed through siRNA screening according to the Human Protein Atlas.^25^

| Gene Name | Expression levels in HeLa (nPTM) | Protein Atlas URL |
| --- | --- | --- |
| Rab1A | 135.5 | https://www.proteinatlas.org/ENSG00000138069-RAB1A |
| Rab1B | 190.9 | https://www.proteinatlas.org/ENSG00000174903-RAB1B |
| Rab2A | 65.7 | https://www.proteinatlas.org/ENSG00000104388-RAB2A |
| Rab2B | 6.1 | https://www.proteinatlas.org/ENSG00000129472-RAB2B |
| Rab3A | 6.3 | https://www.proteinatlas.org/ENSG00000105649-RAB3A |
| Rab3B | 1.1 | https://www.proteinatlas.org/ENSG00000169213-RAB3B |
| Rab3C | 1.7 | https://www.proteinatlas.org/ENSG00000152932-RAB3C |
| Rab3D | 6.6 | https://www.proteinatlas.org/ENSG00000105514-RAB3D |
| Rab4A | 11.1 | https://www.proteinatlas.org/ENSG00000168118-RAB4A |
| Rab4B | 6.0 | https://www.proteinatlas.org/ENSG00000167578-RAB4B |
| Rab5A | 31.5 | https://www.proteinatlas.org/ENSG00000144566-RAB5A |
| Rab5B | 44.5 | https://www.proteinatlas.org/ENSG00000111540-RAB5B |
| Rab5C | 165.4 | https://www.proteinatlas.org/ENSG00000108774-RAB5C |
| Rab6A | 69.4 | https://www.proteinatlas.org/ENSG00000175582-RAB6A |
| Rab6B | 2.4 | https://www.proteinatlas.org/ENSG00000154917-RAB6B |
| Rab6C | Not detected | https://www.proteinatlas.org/ENSG00000222014-RAB6C |
| Rab7A | 100.9 | https://www.proteinatlas.org/ENSG00000075785-RAB7A |
| Rab7B | Not detected | https://www.proteinatlas.org/ENSG00000276600-RAB7B |
| Rab8A | 73.4 | https://www.proteinatlas.org/ENSG00000167461-RAB8A |
| Rab8B | 7.7 | https://www.proteinatlas.org/ENSG00000166128-RAB8B |
| Rab9A | 34.4 | https://www.proteinatlas.org/ENSG00000123595-RAB9A |
| Rab9B | Not detected | https://www.proteinatlas.org/ENSG00000123570-RAB9B |
| Rab10 | 58.6 | https://www.proteinatlas.org/ENSG00000084733-RAB10 |
| Rab11A | 277.9 | https://www.proteinatlas.org/ENSG00000103769-RAB11A |
| Rab11B | 88.2 | https://www.proteinatlas.org/ENSG00000185236-RAB11B |
| Rab12 | 22.1 | https://www.proteinatlas.org/ENSG00000206418-RAB12 |
| Rab13 | 76.6 | https://www.proteinatlas.org/ENSG00000143545-RAB13 |
| Rab14 | 95.8 | https://www.proteinatlas.org/ENSG00000119396-RAB14 |
| Rab15 | 1.2 | https://www.proteinatlas.org/ENSG00000139998-RAB15 |
| Rab17 | 0.2 | https://www.proteinatlas.org/ENSG00000124839-RAB17 |
| Rab18 | 66.7 | https://www.proteinatlas.org/ENSG00000099246-RAB18 |
| Rab19 | 0.3 | https://www.proteinatlas.org/ENSG00000146955-RAB19 |
| Rab20 | 4.8 | https://www.proteinatlas.org/ENSG00000139832-RAB20 |
| Rab21 | 3.7 | https://www.proteinatlas.org/ENSG00000080371-RAB21 |
| Rab22A | 10.4 | https://www.proteinatlas.org/ENSG00000124209-RAB22A |
| Rab23 | 11.6 | https://www.proteinatlas.org/ENSG00000112210-RAB23 |
| Rab24 | 20.3 | https://www.proteinatlas.org/ENSG00000169228-RAB24 |
| Rab25 | Not detected | https://www.proteinatlas.org/ENSG00000132698-RAB25 |
| Rab26 | 12.3 | https://www.proteinatlas.org/ENSG00000167964-RAB26 |
| Rab27A | 30.4 | https://www.proteinatlas.org/ENSG00000069974-RAB27A |
| Rab27B | 3.5 | https://www.proteinatlas.org/ENSG00000041353-RAB27B |
| Rab28 | 19.1 | https://www.proteinatlas.org/ENSG00000157869-RAB28 |
| Rab7L1 (Rab29) | 10.3 | https://www.proteinatlas.org/ENSG00000117280-RAB29 |
| Rab30 | 16.9 | https://www.proteinatlas.org/ENSG00000137502-RAB30 |
| Rab31 | 116.8 | https://www.proteinatlas.org/ENSG00000168461-RAB31 |
| Rab32 | 132.3 | https://www.proteinatlas.org/ENSG00000118508-RAB32 |
| Rab33A | 0.3 | https://www.proteinatlas.org/ENSG00000134594-RAB33A |
| Rab33B | 4.1 | https://www.proteinatlas.org/ENSG00000172007-RAB33B |
| Rab34 | 154.4 | https://www.proteinatlas.org/ENSG00000109113-RAB34 |
| Rab35 | 36.1 | https://www.proteinatlas.org/ENSG00000111737-RAB35 |
| Rab36 | 0.3 | https://www.proteinatlas.org/ENSG00000100228-RAB36 |
| Rab37 | 0.1 | https://www.proteinatlas.org/ENSG00000172794-RAB37 |
| Rab38 | 5.3 | https://www.proteinatlas.org/ENSG00000123892-RAB38 |
| Rab39 | Not detected | https://www.proteinatlas.org/ENSG00000179331-RAB39A |
| Rab39B | Not detected | https://www.proteinatlas.org/ENSG00000155961-RAB39B |
| Rab40A | 1.7 | https://www.proteinatlas.org/ENSG00000172476-RAB40A |
| Rab40B | 10.9 | https://www.proteinatlas.org/ENSG00000141542-RAB40B |
| Rab40C | 12.3 | https://www.proteinatlas.org/ENSG00000197562-RAB40C |
| Rab41 | 0.2 | https://www.proteinatlas.org/ENSG00000147127-RAB41 |
| Rab42 | 7.6 | https://www.proteinatlas.org/ENSG00000188060-RAB42 |
| Rab43 | 2.6 | https://www.proteinatlas.org/ENSG00000172780-RAB43 |

**Supplementary Figure 1:**
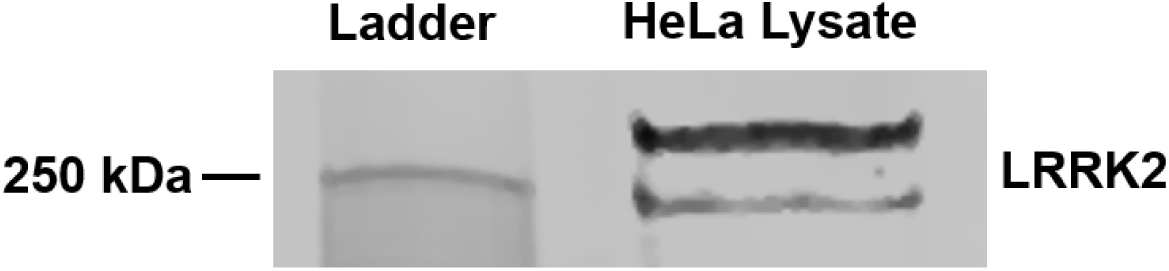
LRRK2 protein is expressed in mt-mKeima/YFP-Parkin HeLa cells. Western immunoblot of untreated mt-mKeima/YFP-Parkin HeLa cell lysates was probed with rabbit anti-LRRK2 to show endogenous LRRK2 expression.

**Supplementary Table 2:**
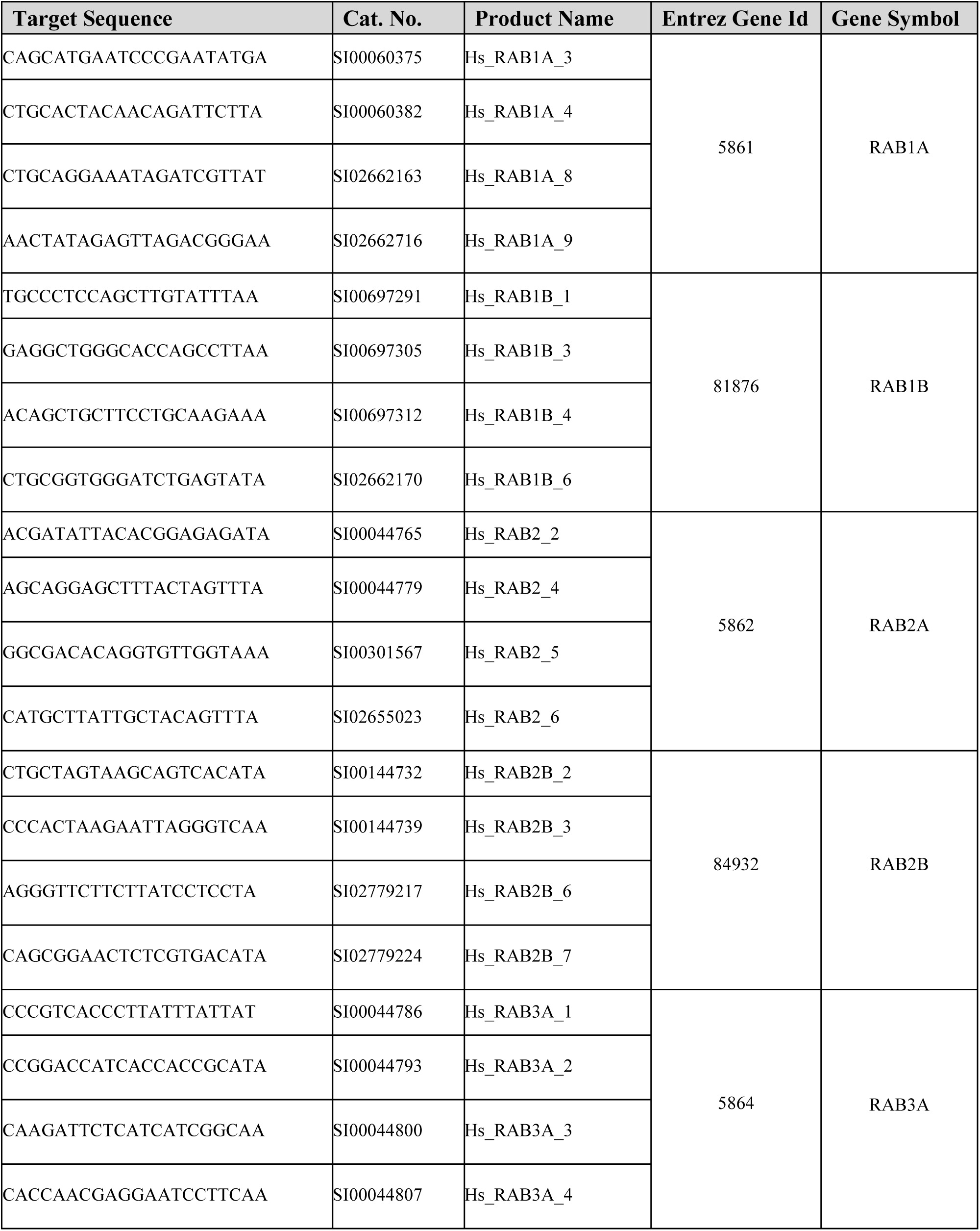
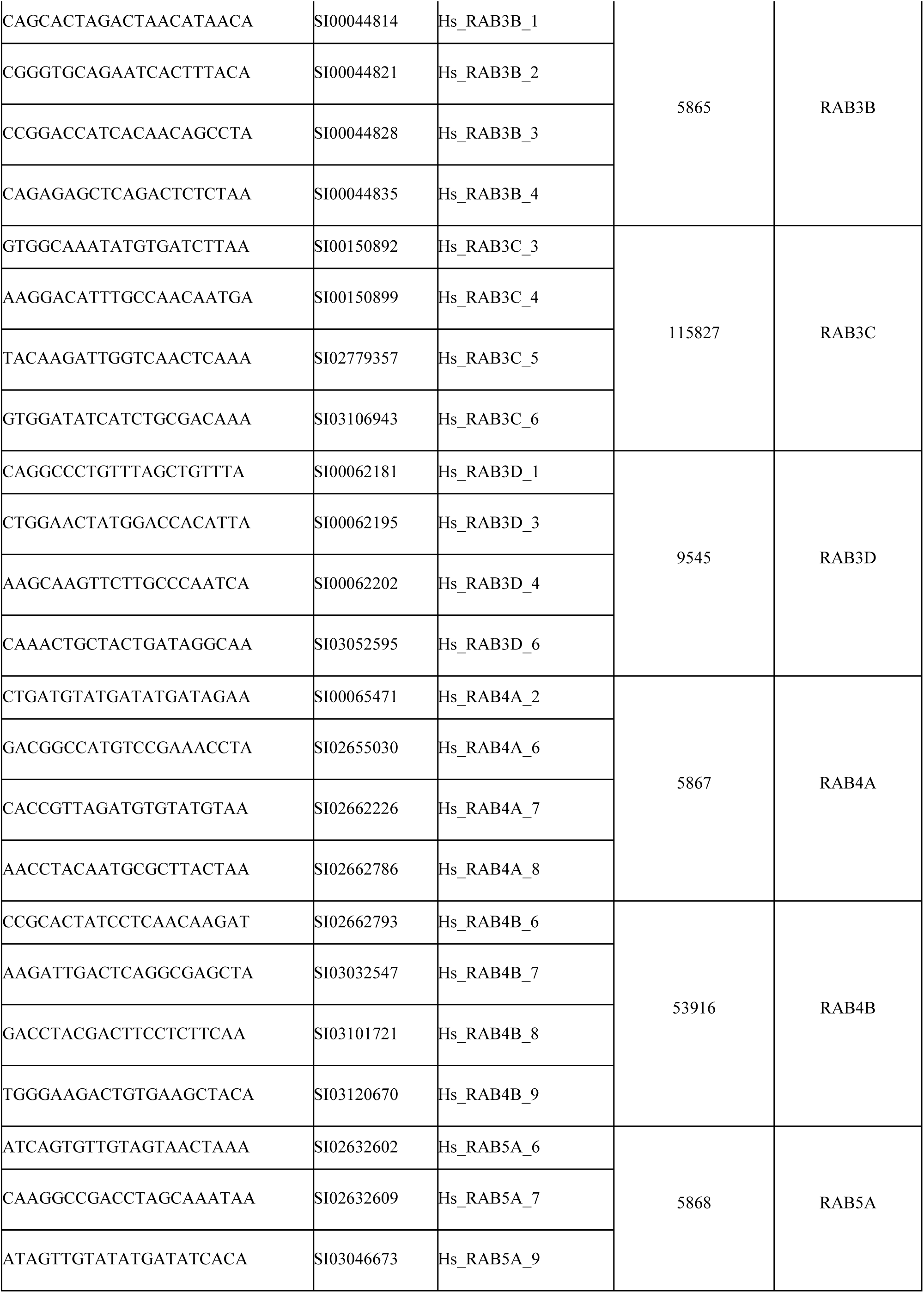
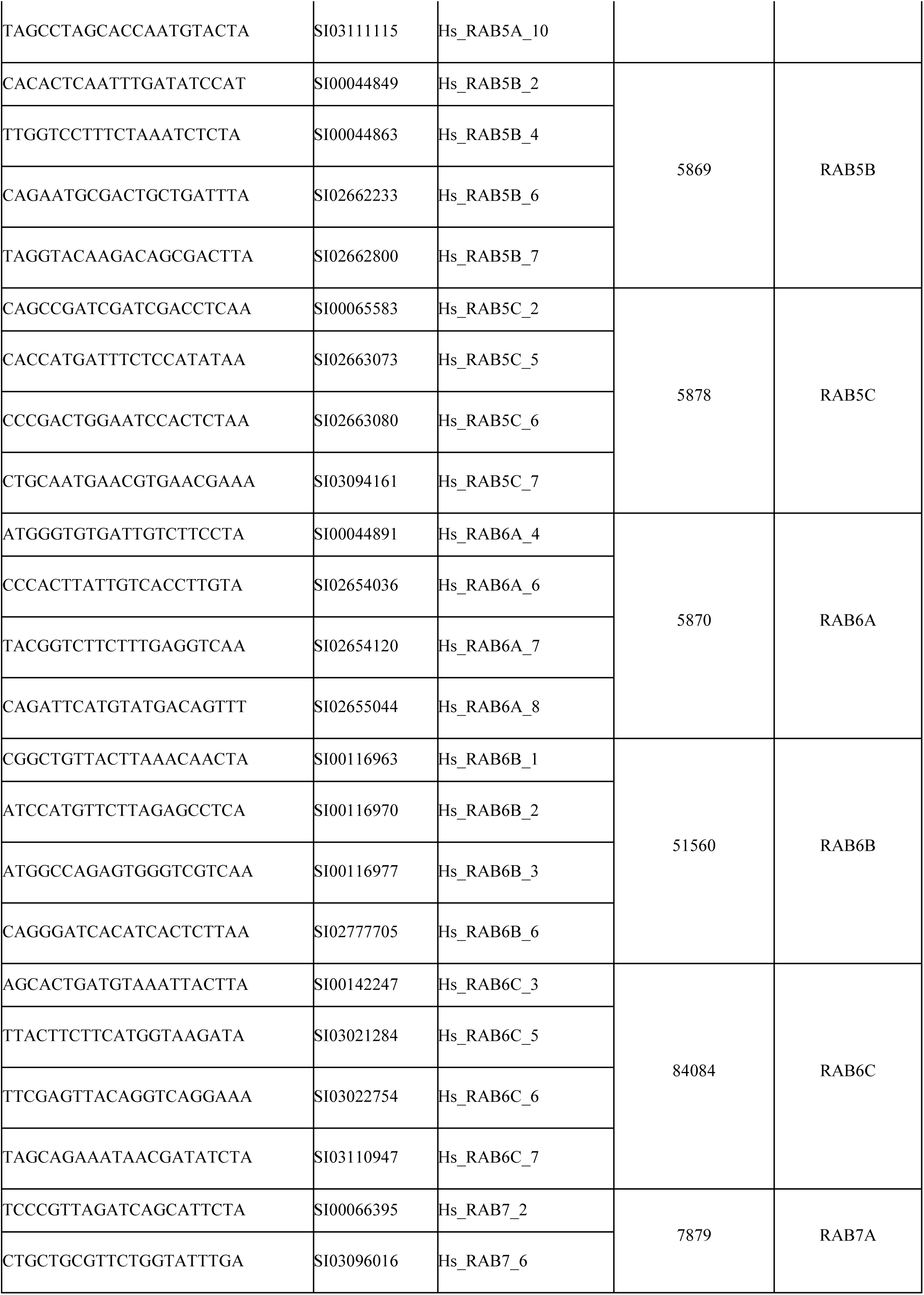
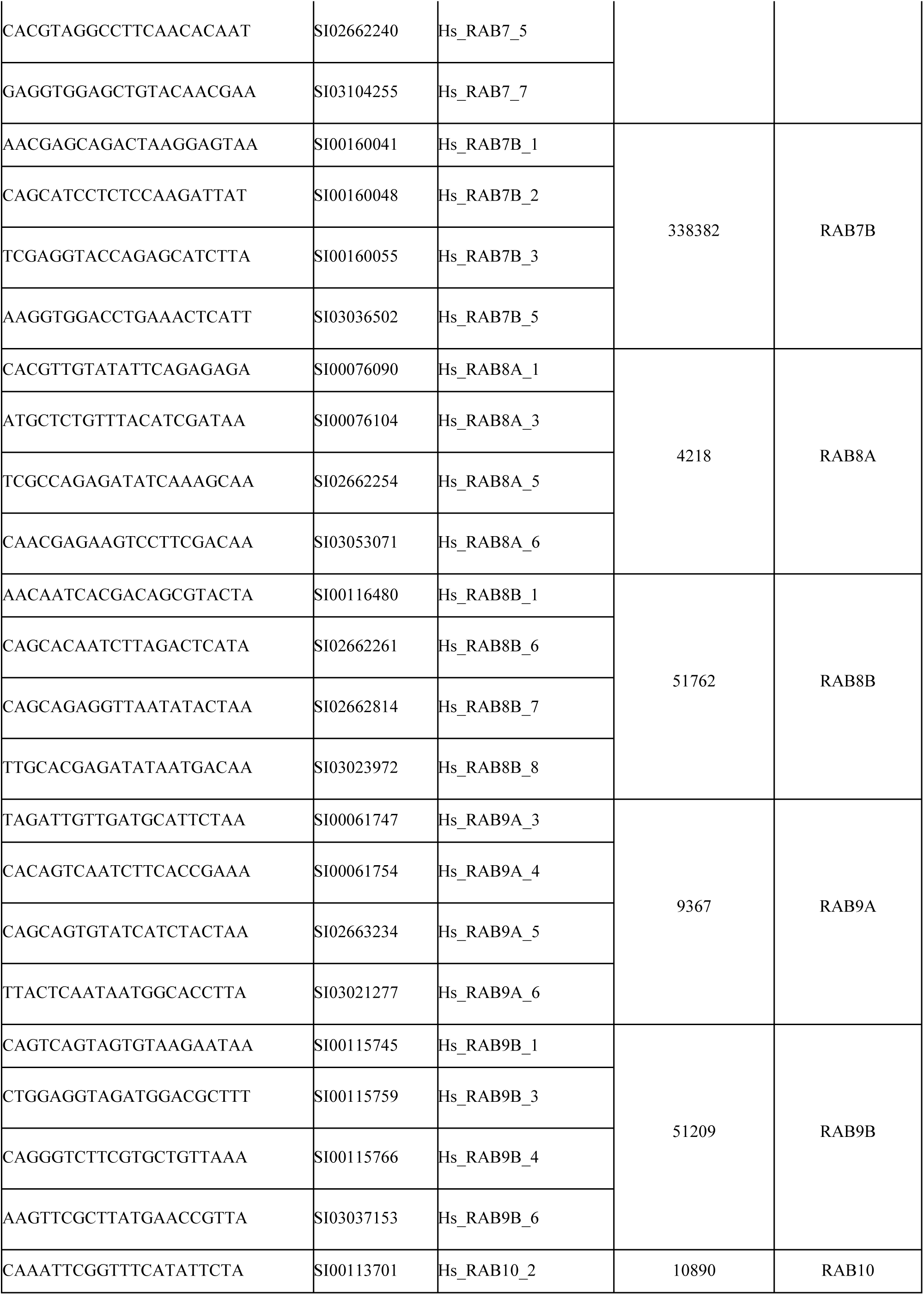
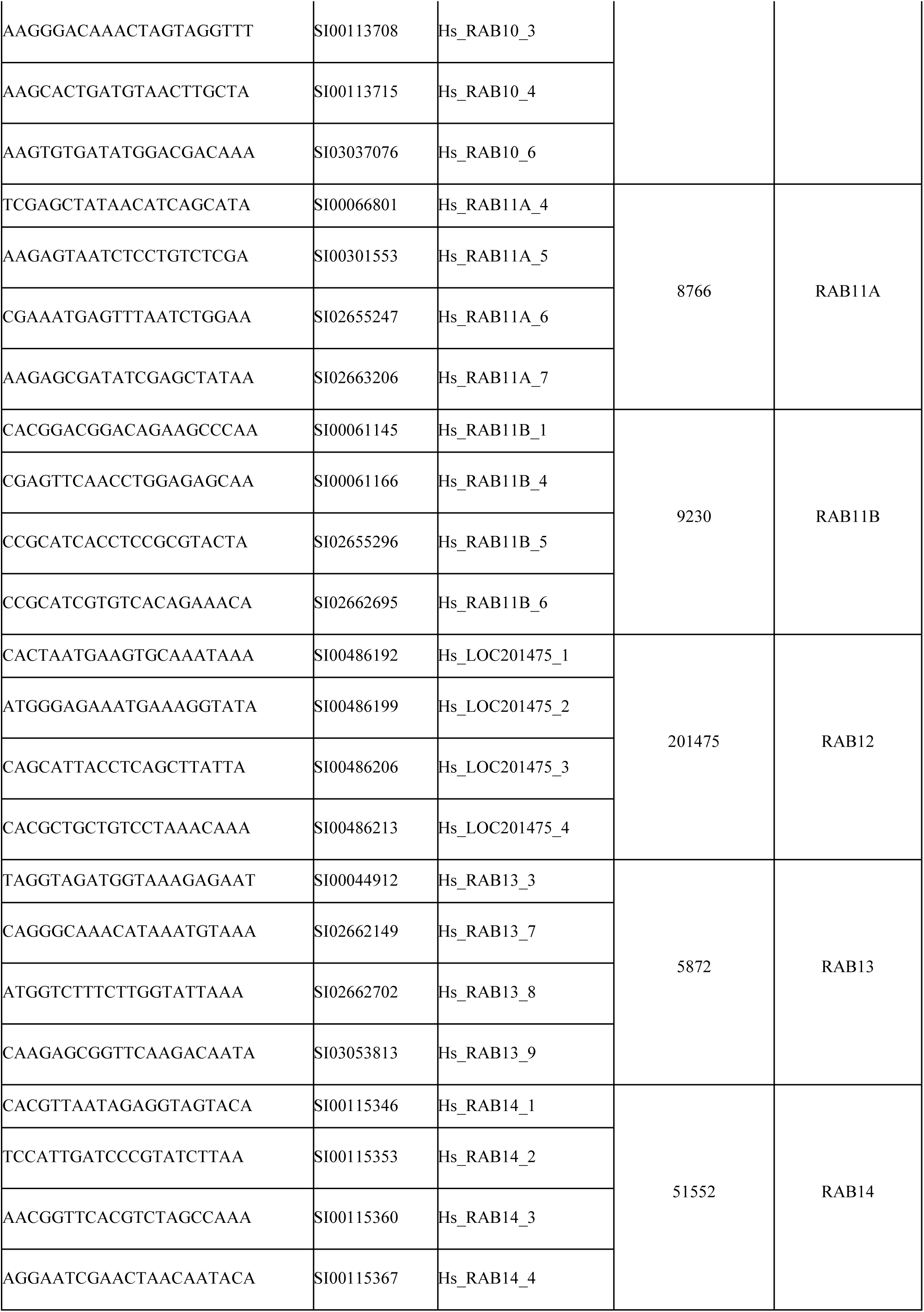
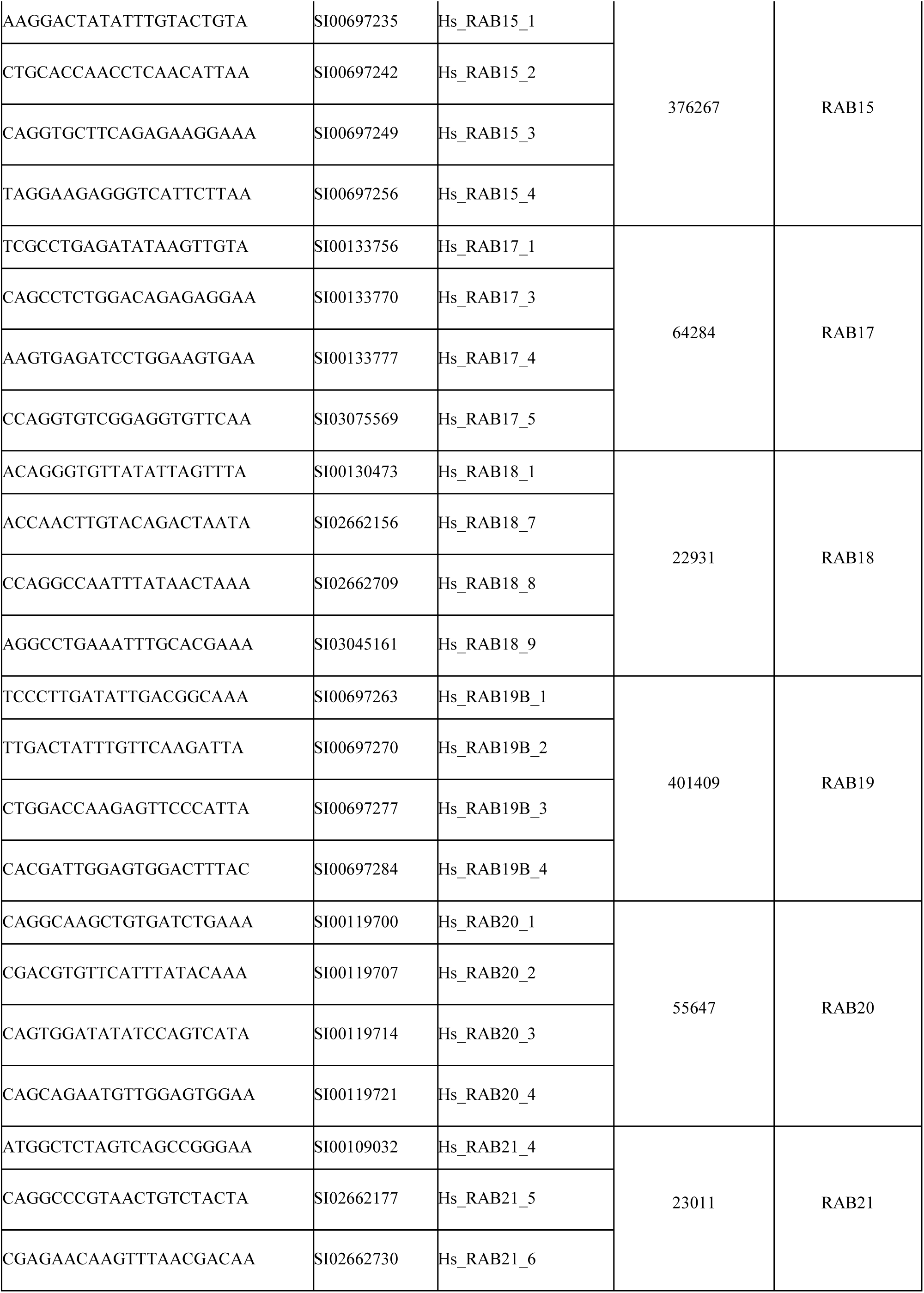
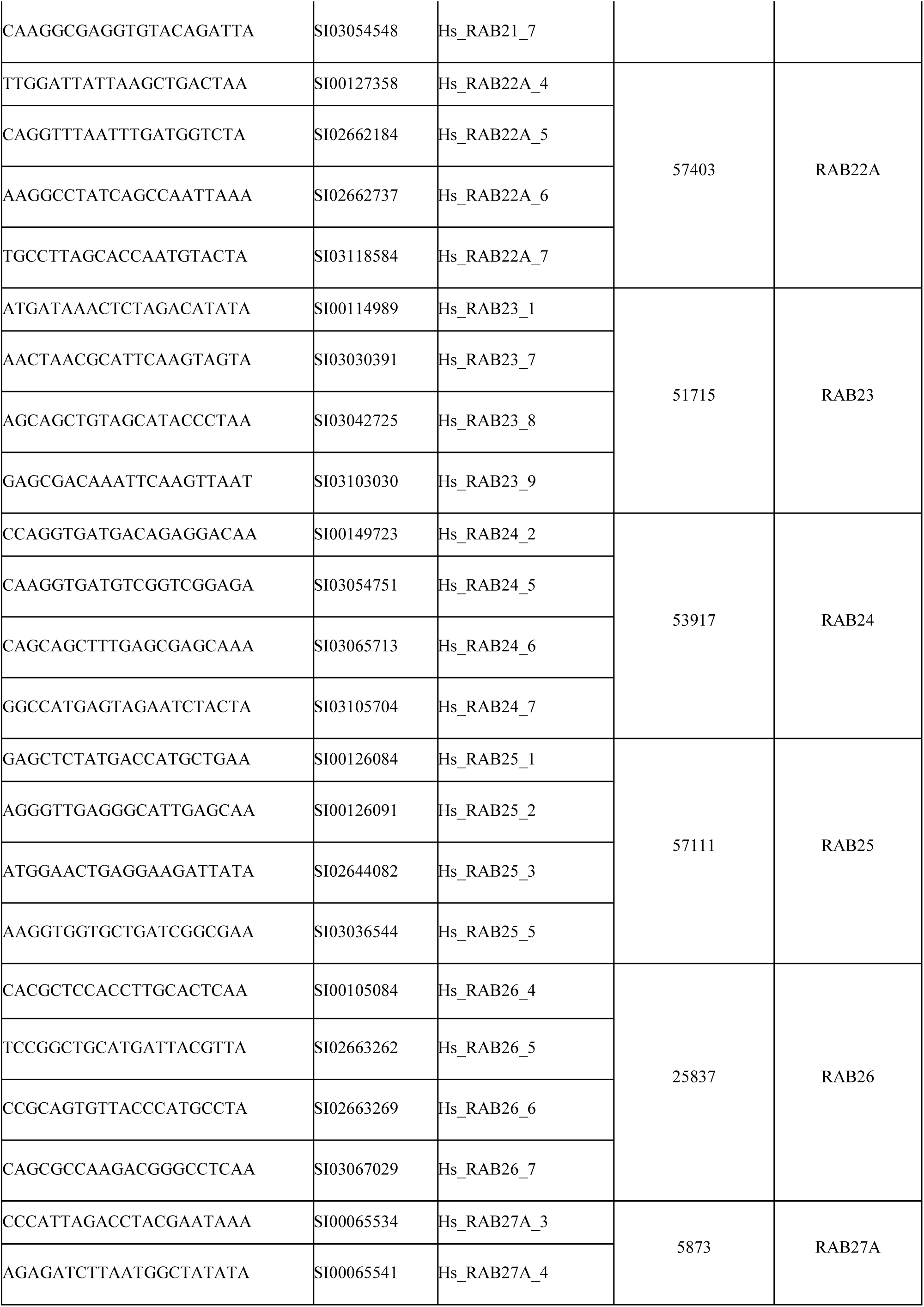
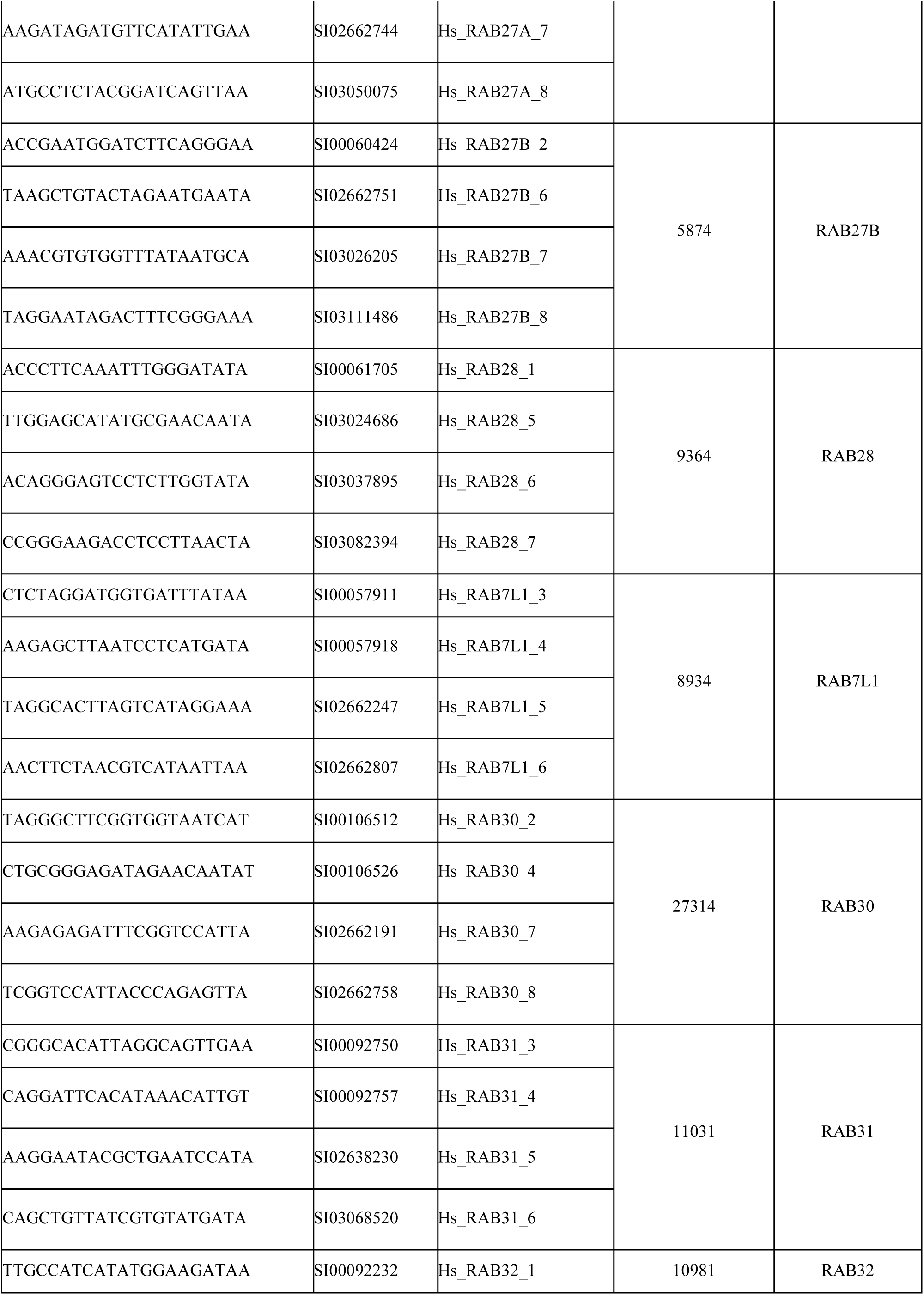
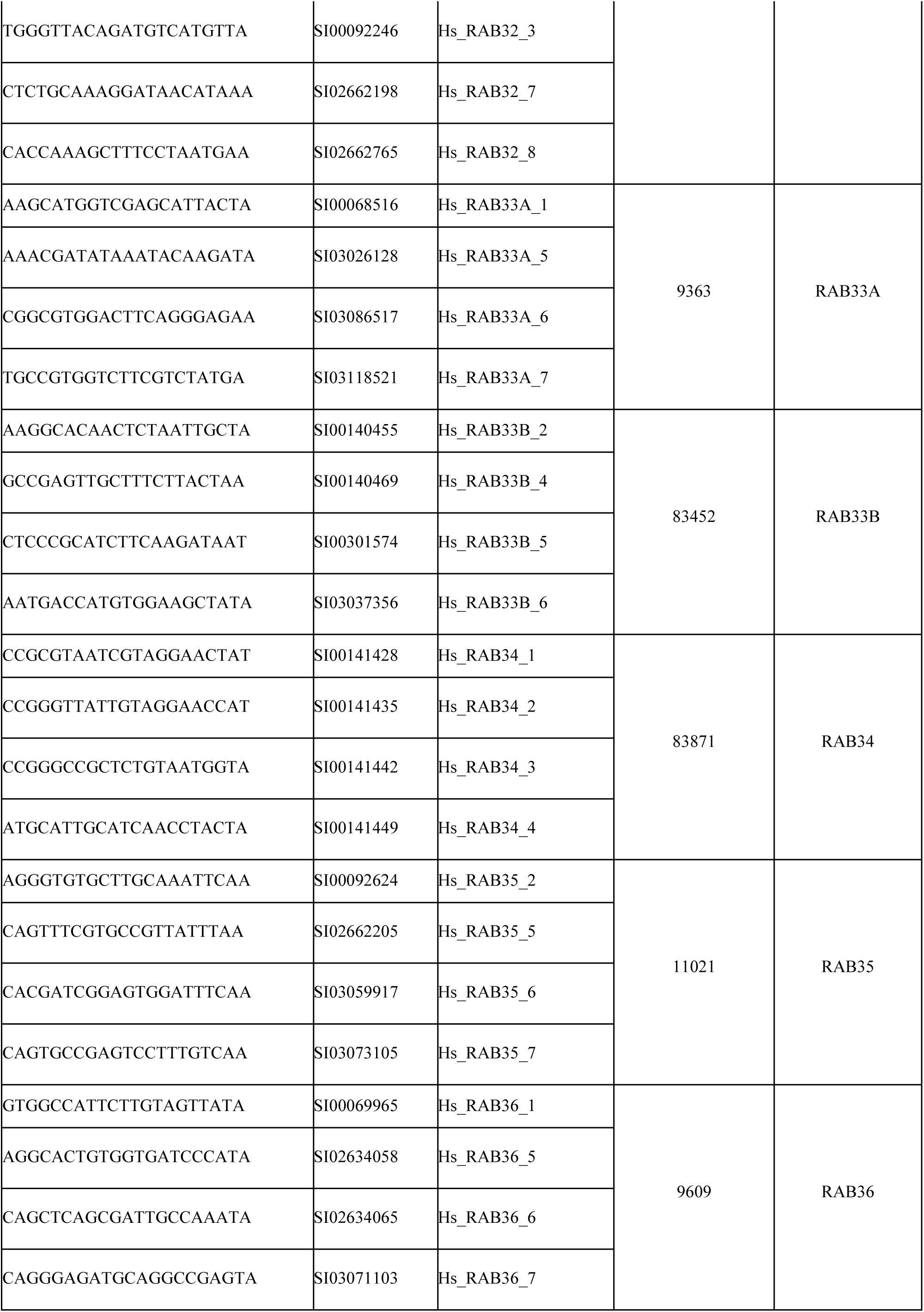
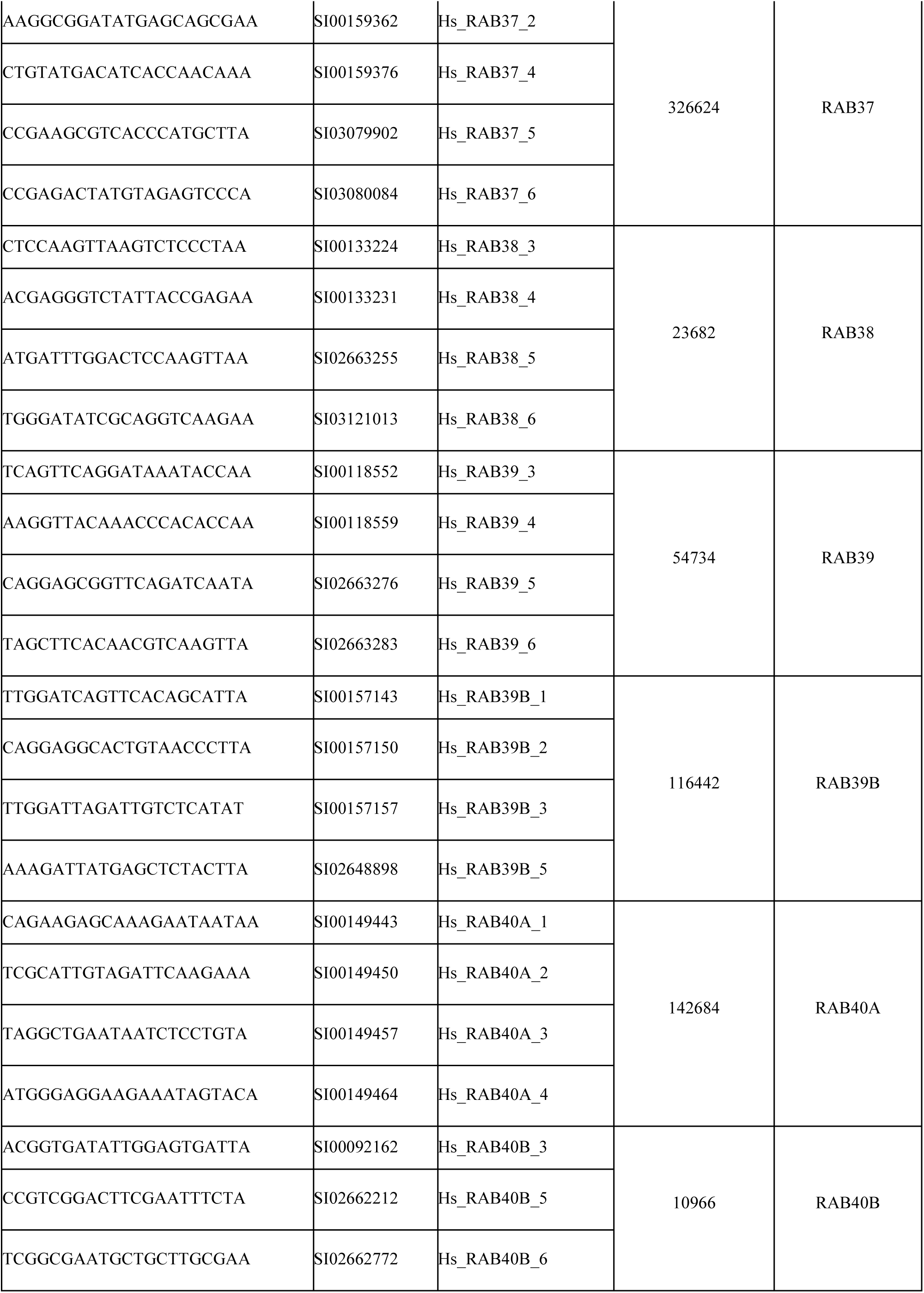
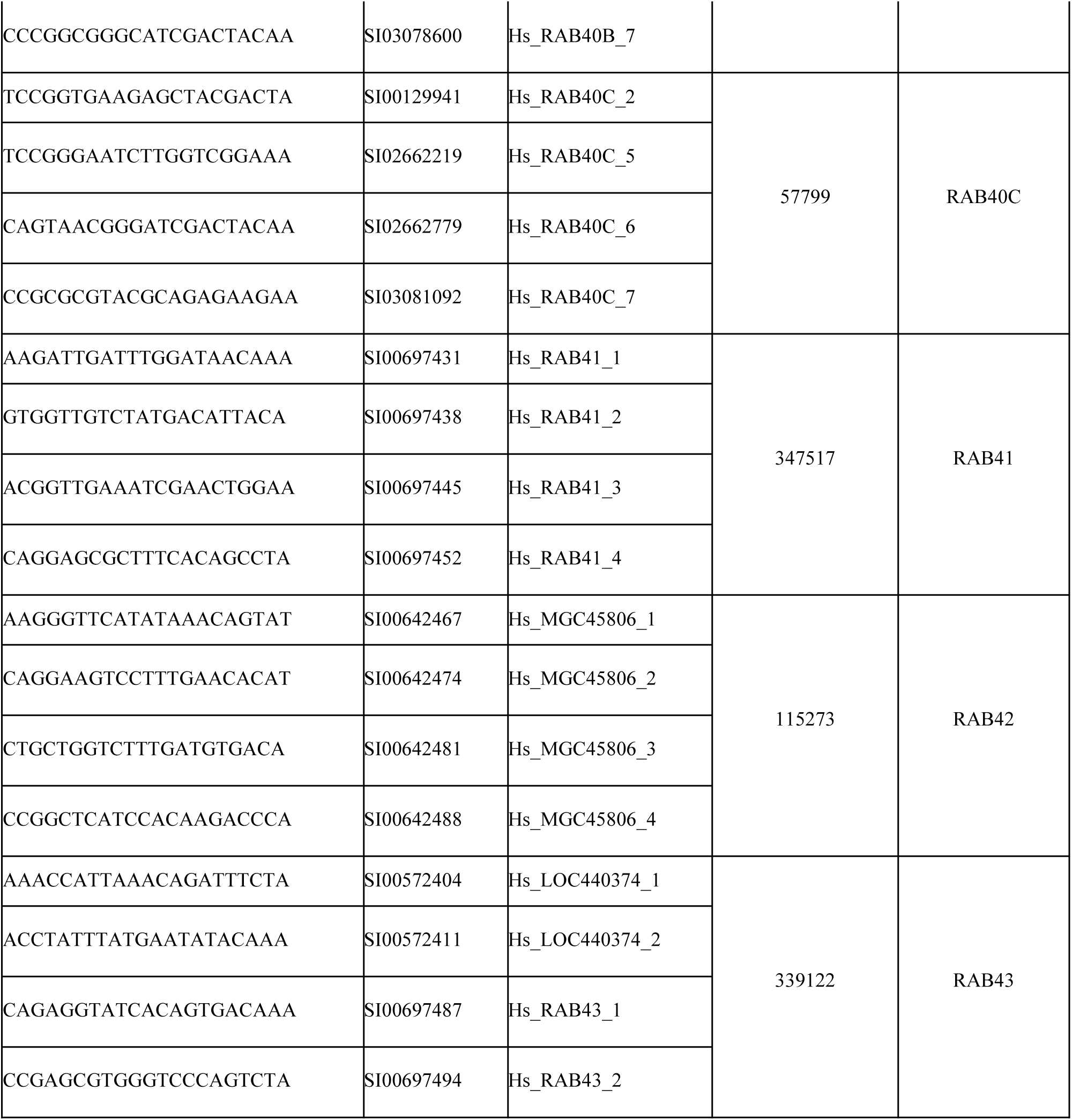
Sequence and catalogue information of Rab family siRNA library. Sequences, Qiagen catalog numbers, and gene identifier information for the siRNA library against human Rab family proteins utilized in mitophagy screening.

**Supplementary Figure 2:**
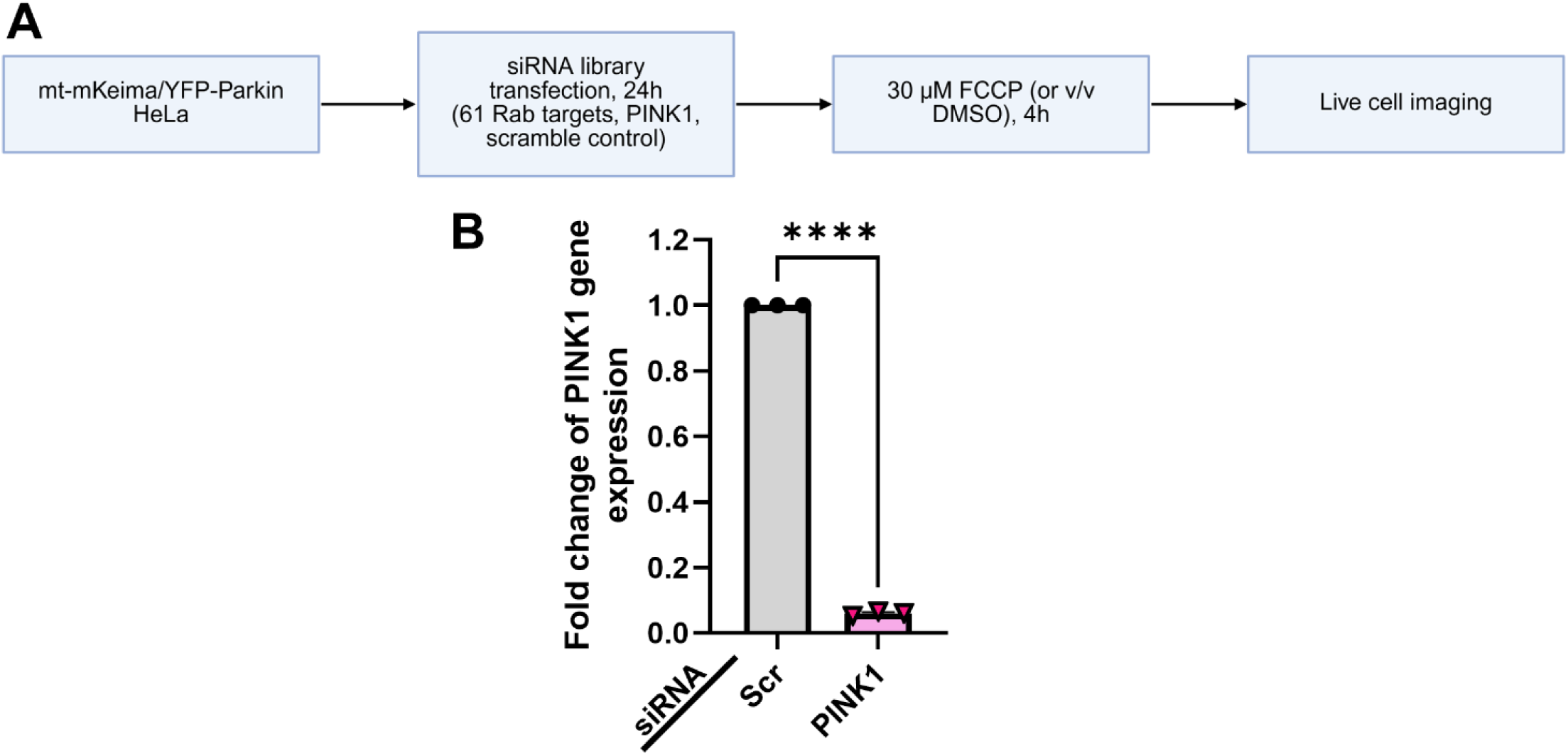
Rab protein siRNA mitophagy screen protocol workflow and validation. (**A**) Outline of Rab family siRNA mitophagy screening protocol. HeLa cells stably expressing mt-mKeima and YFP-Parkin were plated onto 96-well plates pre-stamped with an siRNA library spanning the Rab family of proteins, along with a scramble negative control siRNA and a PINK1 siRNA as a positive control for mitophagy impairment. Following 24 h of transfection, cells were treated with FCCP (or an equivalent volume of DMSO vehicle control) for 4 h to induce mitophagy, followed by live-cell imaging for mitophagy analysis. Figure generated with BioRender. (**B**) RNA was collected for qRT-PCR analysis of PINK1 mRNA expression in mt-mKeima/YFP-parkin HeLa cells following reverse transfection with siRNA against PINK1 (siPINK1) or the scramble control (siScr) for 24 h. Internal control for normalization was GAPDH. \*\*\*\**p* < 0.0001, as determined by unpaired t-test. *n* = 3 biological replicates. Data are presented as mean ± SEM.

**Supplementary Figure 3:**
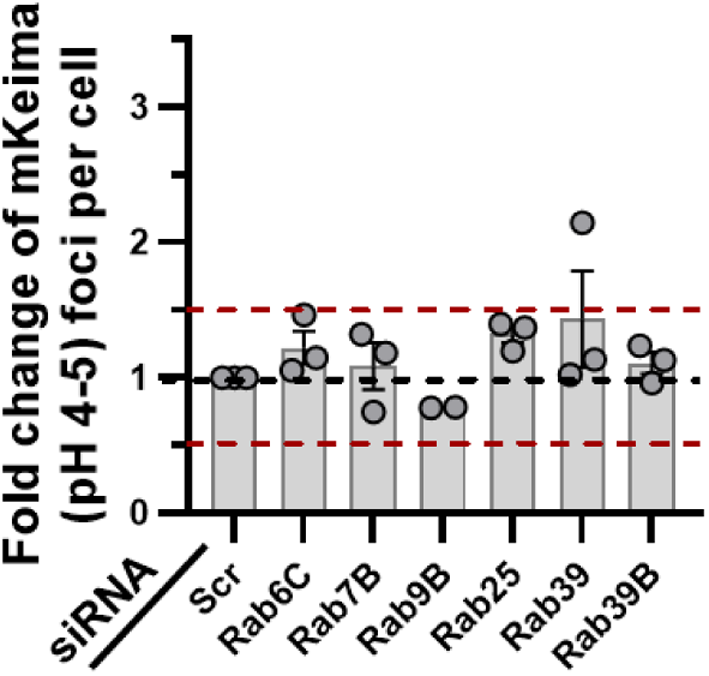
siRNA against Rab proteins not expressed in HeLa cells did not alter mitophagy based on thresholding parameters selected for the siRNA screen. Endogenous expression levels of Rab family proteins were collected from the Human Protein Atlas. Candidates in the Rab family siRNA library utilized in mitophagy screening with no endogenous expression in HeLa cells were separated from candidates expressed in HeLa (Figure 2) and instead are displayed here. siRNAs targeting these non-expressed candidates had no impact on mitophagy based on the selected cutoff of a 50% increase or decrease in fold change of mt-mKeima (pH 4-5) foci per cell. *n =* 3 replicates (except Rab9B: *n =* 2 replicates).

**Supplementary Figure 4:**
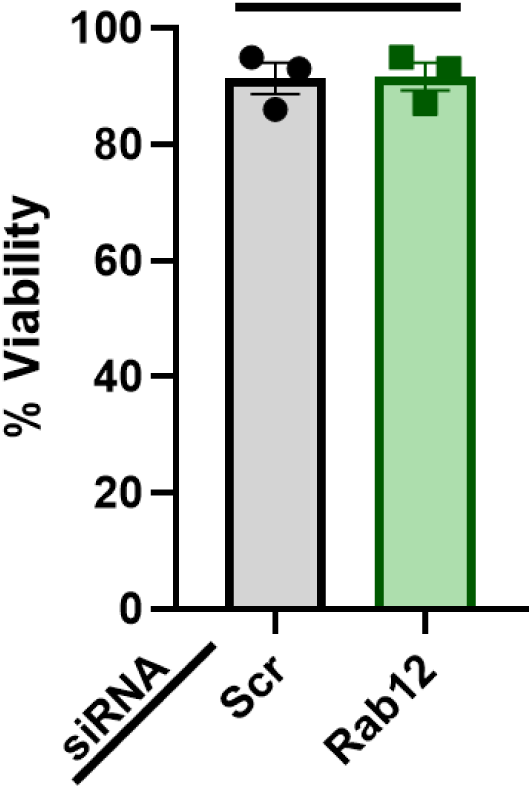
Cell viability analysis in mt-mKeima/YFP-Parkin HeLa cells. The trypan blue exclusion assay was used to quantify percent viability in mt-mKeima/YFP-Parkin HeLa cultures transfected with scramble (Scr) siRNA or siRNA against Rab12 for 72 h. No difference in viability was observed with Rab12 knockdown relative to siScr transfection control. Data was analyzed using an unpaired t-test. *n* = 3 biological replicates. Data are presented as mean ± SEM.

**Supplementary Figure 5:**
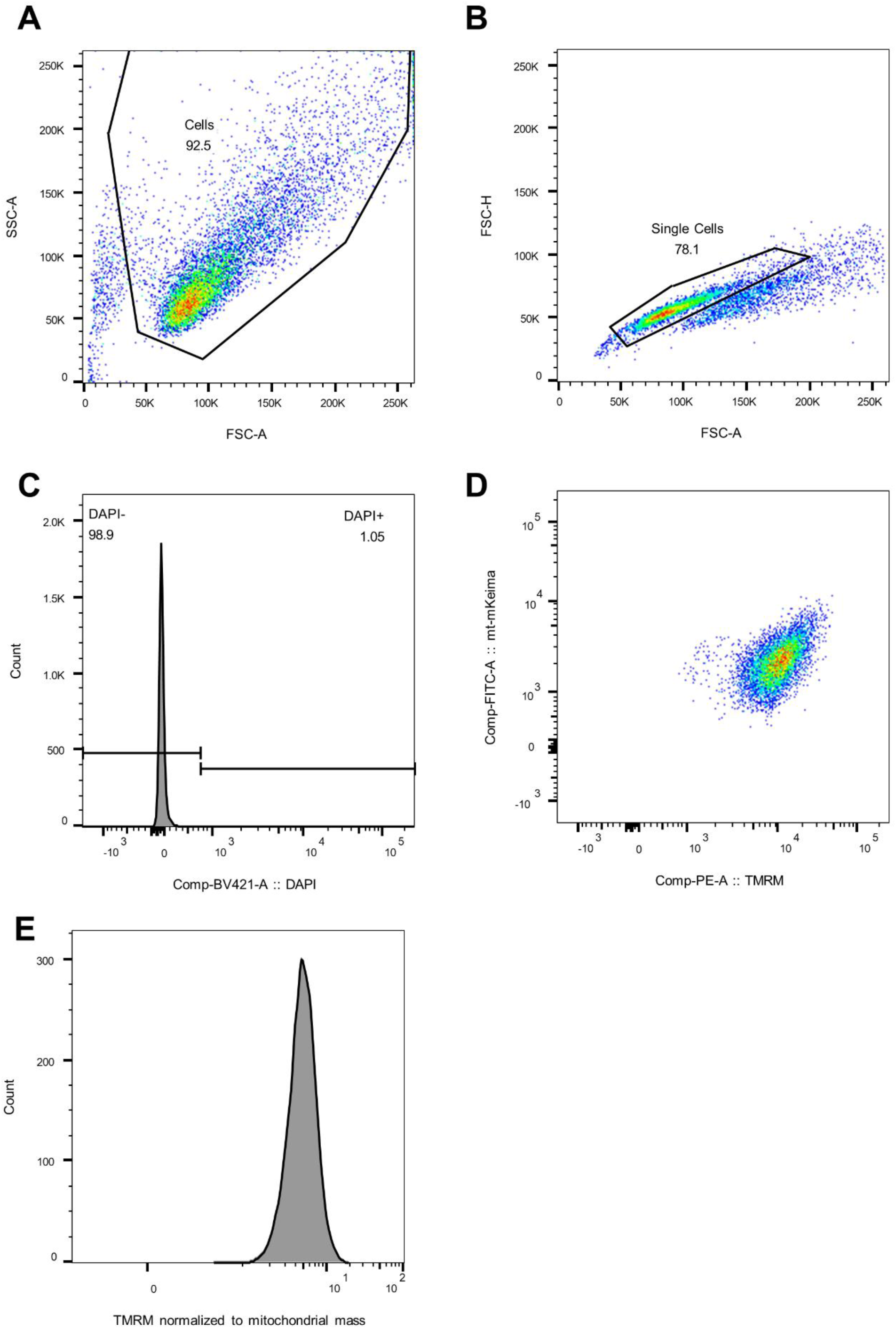
Sample of siScr HeLa cells demonstrating gating strategy for flow cytometry after applying the compensation matrix. **(A**) FSC/SSC gated to exclude debris. (**B**) Single cells gated in plot of FSC-A/FSC-H, excluding doublets and aggregates. (**C**) Dead cells excluded by selecting events negative for DAPI. (**D**) Parameters of interest, TMRM signal versus mt-mKeima. (**E**) Histogram of the derived parameter - TMRM signal divided by mt-mKeima - representing TMRM normalized to mitochondrial mass per cell.

**Supplementary Figure 6:**
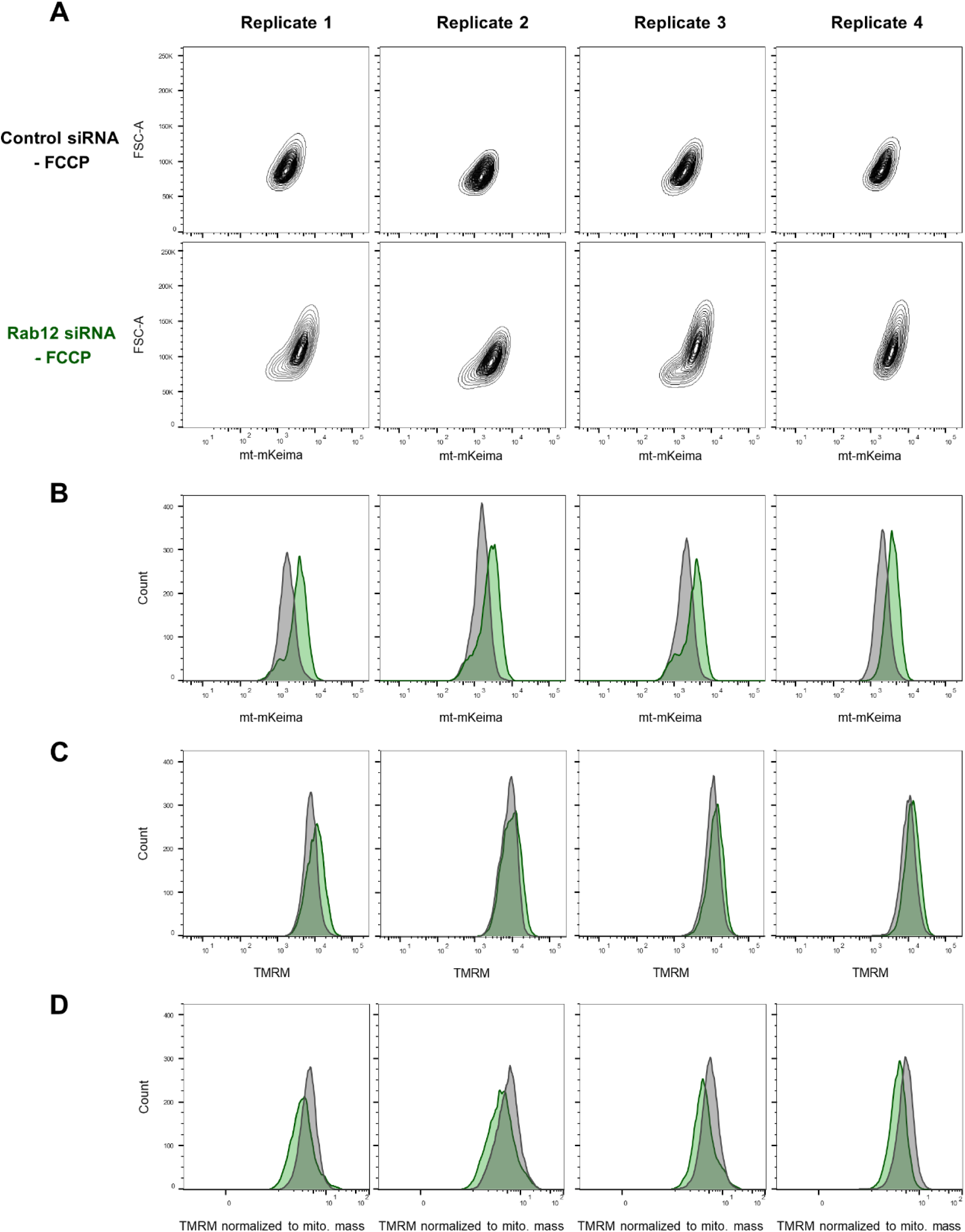
Plots of siScr HeLa cells (gray) compared siRab12 HeLa cells (green) prior to FCCP exposure for four independent experiments. (**A**) mt-mKeima signal versus forward scatter (FSC). (**B**) Histograms of mt-mKeima signal. (**C**) Histograms of TMRM signal. (**D**) Histograms of the derived parameter, mt-mKeima divided by TMRM signal.

**Supplementary Figure 7:**
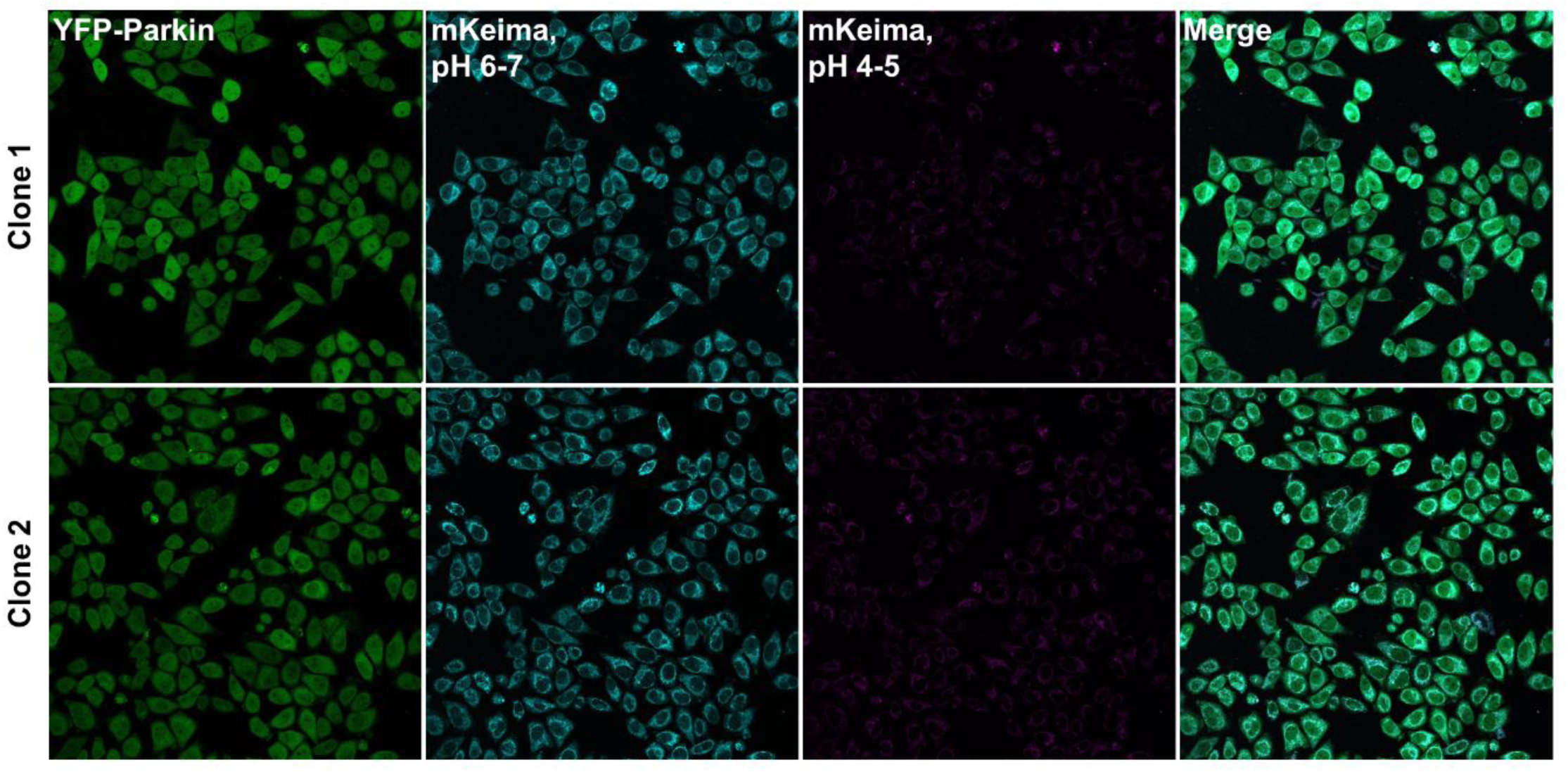
Comparison of mt-mKeima/YFP-Parkin clones utilized in siRNA screening (Clone 1) and validation of Rab12’s impact on mitochondrial homeostatic mechanisms (Clone 2). Cultures of mt-mKeima/YFP-Parkin HeLa cells exhibiting heterogeneous fluorescence levels of mt-mKeima and YFP-Parkin were single-cell sorted, and two separate clones were selected for siRNA screening (Clone 1) and validation of Rab12 impact on mitophagy (Clone 2). Representative 20X confocal images of the selected clones presented here display YFP-Parkin (green), mt-mKeima at neutral pH (cyan), and mt-mKeima at lysosomal pH (magenta), and were taken with identical laser settings to show similar fluorescence levels of both mt-mKeima and YFP-Parkin protein in each clone.

